# Topology-driven classification of time series

**DOI:** 10.64898/2026.04.25.720787

**Authors:** Alexandra Bernadotte

## Abstract

Time series analysis is fundamentally limited by the lack of representations that reflect the underlying generative mechanisms of observed signals. Existing approaches, ranging from spectral decompositions to modern machine learning, primarily operate on signal values or frequency content, and therefore fail to capture the intrinsic structure of the dynamics that produce the data.

In this work, we introduce a geometric framework that establishes a direct correspondence between the generative structure of a time series and the topology of its delay embedding. We show that broad classes of signals (including exponential, harmonic, and exponentially modulated oscillatory processes) induce invariant low-dimensional subspaces in Hankel embedding space, which dimension is determined solely by the number and type of latent dynamical components.

This leads to a unifying principle: the intrinsic dimension and geometry of delay embeddings act as invariants of the underlying dynamics. Building on this result, we reformulate time series classification as the problem of separating equivalence classes defined by *ε*-neighborhoods of subspaces on a Grassmann manifold. This yields a topological classifier that is interpretable, data-efficient, and provably robust, where noise admits a natural geometric interpretation as bounded perturbations of subspaces.

We demonstrate that the proposed framework distinguishes signals with indistinguishable spectral signatures and consistently recovers the latent structure of complex, noisy, multi-component processes. On benchmark EEG data, the method achieves state-of-the-art performance without feature engineering or large-scale training.

These results suggest a shift from feature-based and statistical representations toward a geometric theory of time series, in which structure, classification are governed by the topology of embeddings.

An interactive web-based demonstration is available to facilitate exploration of the geometric structure of delay embeddings and the proposed classification approach.

## Introduction

A central challenge in time series analysis is to infer the underlying dynamical structure from observed data. Classical approaches, including Fourier and wavelet analysis, represent signals as superpositions of predefined basis functions but do not explicitly encode the mechanisms that generate them. As a consequence, signals produced by fundamentally different processes may exhibit nearly identical spectral characteristics, limiting the discriminative power of frequency-based representations.

An alternative perspective is provided by delay embeddings, where a time series is mapped into a higher-dimensional space that captures its temporal structure. While such embeddings have been widely used in dynamical systems theory and singular spectrum analysis, their geometric properties have not been fully exploited for classification.

In this work, we show that the geometry of delay embeddings provides a complete and interpretable representation of a broad class of dynamical systems. Specifically, we establish that signals generated by exponential, harmonic, and exponentially modulated oscillatory processes correspond to trajectories confined to low-dimensional linear subspaces, which dimension is determined by the number and type of latent components.

This observation leads to a fundamental reformulation of time series classification: instead of operating on signal values or spectral features, classification can be posed as the problem of identifying the subspace (and its *ε* −neighborhood) in which the embedded trajectory resides. In this sense, the topology of the embedding becomes a direct observable of the underlying dynamics.

This geometric viewpoint yields a classifier that is both interpretable and robust. Unlike data-driven models, it does not require large training datasets; unlike spectral methods, it captures structural differences between signals with similar frequency content. Moreover, noise admits a natural geometric interpretation as a perturbation that induces an *ε* −neighborhood around the underlying subspace, providing theoretical guarantees of stability.

Together, these results establish a general geometric paradigm for time series analysis, linking generative structure, intrinsic dimension, and classification within a unified framework.

From this perspective, time series classification can be interpreted as a problem of geometric inference in embedding space.

## Related Work

Time series classification has been extensively studied across several paradigms, including spectral analysis, statistical feature extraction, and modern machine learning [1, 2]. Despite substantial progress, these approaches typically operate on representations that do not directly reflect the underlying generative structure of the signal. In this section, we review the most relevant directions and position the proposed method within this landscape.

### Spectral and decomposition-based methods

Classical time series analysis relies on spectral representations such as Fourier and wavelet transforms [3, 4]. These methods decompose signals into predefined basis functions and are effective for stationary or weakly non-stationary processes. However, they do not explicitly encode the geometry of the underlying dynamics. As a result, signals generated by fundamentally different mechanisms may exhibit similar spectral signatures, limiting the discriminative power of frequency-based representations.

Closely related are embedding-based approaches, including singular spectrum analysis (SSA) and subspace identification [5, 6, 7]. These methods exploit low-rank structure in Hankel embeddings and establish connections between time series and Grassmannian geometry [7]. While they reveal that structured signals admit low-dimensional representations, their primary use has been in decomposition, denoising, and forecasting, rather than classification.

Recent work has extended Hankel-based approaches to tasks such as estimating the number of latent sources and real-time signal processing pipelines [8, 9, 10]. However, these methods do not explicitly leverage embedding geometry for classification.

### Statistical and feature-based methods

A large class of methods relies on handcrafted features, including statistical moments, autocorrelation measures, and shape descriptors [1]. These features are subsequently used with standard classifiers such as support vector machines or random forests. Although effective in many applications, such approaches are inherently dependent on feature design and may fail to generalize across signals with different generative mechanisms.

### Machine learning and deep learning approaches

Recent advances in time series classification are driven by deep learning architectures, including convolutional, recurrent, and transformer-based models [11, 12]. These methods can achieve high predictive accuracy when large labeled datasets are available. However, they typically require extensive training, offer limited interpretability, and do not explicitly capture the geometric or dynamical structure of the signal.

### Topological data analysis

Topological data analysis (TDA) provides tools for capturing the global shape of data, most notably through persistent homology [13, 14]. In time series analysis, TDA is often applied to delay embeddings, yielding persistence diagrams or related summaries for classification [15]. While these methods provide robustness to noise and capture global topological features, they rely on secondary representations that must be processed by downstream learning models [16]. Moreover, they do not explicitly exploit the linear subspace structure inherent in many classes of signals.

### Subspace and manifold-based methods

Subspace and manifold learning methods aim to represent data as lying near low-dimensional structures embedded in high-dimensional spaces [17, 18]. These approaches are closely related to the present work in their use of geometric representations. However, they are typically designed for clustering or dimensionality reduction, rather than for constructing explicit classification rules based on subspace geometry and tolerance-based separation.

### Position of the present work

The proposed method synthesizes ideas from embedding theory, subspace methods, and topological analysis, while introducing a fundamentally different perspective on classification. In contrast to spectral and feature-based approaches, it operates directly on the geometry of delay embeddings. Unlike deep learning models, it does not require large-scale training and remains interpretable. Compared to classical TDA, it avoids intermediate topological summaries and instead directly exploits the subspace structure of embedded trajectories.

Building on the interpretation of Hankel embeddings as points on Grassmann manifolds [7], we extend this framework from representation to classification. In particular, we formulate time series classification as the problem of separating equivalence classes defined by *ε*-neighborhoods of low-dimensional subspaces. To the best of our knowledge, this formulation—combining subspace geometry, intrinsic dimension, and adaptive tolerance control—has not been previously proposed in the context of time series analysis.

## Methods

### Problem formulation

Let *f* = (*f*_1_, …, *f*_*N*_) be a real-valued time series. The goal is to assign *f* to a class *c* ∈ 𝒞 corresponding to a particular generative mechanism.

We assume that each class induces a low-dimensional linear subspace *S*_*c*_ ⊂ ℝ^*n*^ in the space of delay embeddings (see bellow), and that observed signals (in a form of time series) are perturbed realizations lying within an *ε*-neighborhood of these subspaces.

Formally, classification is defined as

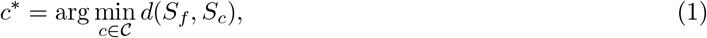

subject to the separability condition

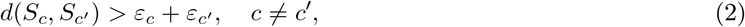

where *S*_*f*_ is the subspace (based on a piecewise linear trajectory *L*_*f*_) associated with the embedding of *f*, and *d*(·, ·) denotes a distance on the Grassmann manifold.

#### 0.0.1 Delay embedding and subspace construction

We interpret time series as trajectories constrained by latent dynamical operators, which action induces invariant subspaces in embedding space. We construct the Hankel (trajectory) matrix on time series *f*

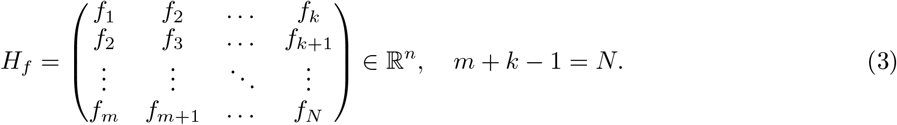

We interpret *H*_*f*_ as a piecewise linear trajectory *L*_*f*_ (columns are dots of *L*_*f*_) in the space of matrices ℝ^*m×k*^ and consider the linear subspace 𝒮_*f*_ spanned by its columns:

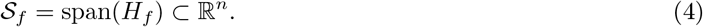

Thus, each time series is associated with a subspace 𝒮_*f*_ . The associated subspace 𝒮_*f*_ is estimated via singular value decomposition (SVD). The intrinsic dimension of *L*_*f*_ is determined by rank estimation of *H*_*f*_, consistent with the theoretical results.

### Theoretical foundation

Structured time series admit low-rank embeddings. In particular, consider signals of the form

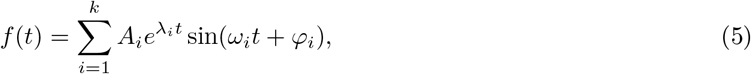

including exponentials, harmonic signals, and exponentially modulated oscillations.

We analyze the topology of piecewise linear trajectories corresponding to time series generated by functions of different types.

#### 0.0.2 Sum of exponential components

Following the approach in [7], we formulate a result concerning the dimension of the space in which the piecewise linear trajectory of a time series lies when the signal is representable as a sum of *k* exponentials of the form Equation 6.

##### Lemma 1

(Rank of exponential signals). Let

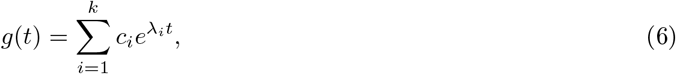

where *c*_*i*_, *λ*_*i*_ are constants with *λ*_*i*_≠ *λ*_*j*_ for *i*≠ *j*. Then the Hankel matrix *H*_*g*(*t*)_ has rank

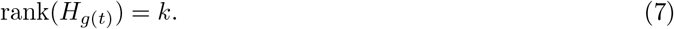

Moreover, the piecewise linear trajectory *K*_*g*(*t*)_ lies in a *k*-dimensional subspace *L* ⊂ ℝ^*n*^, spanned by the vectors

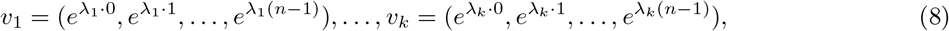

Proof▷ Each exponential component generates a linearly independent sequence. The Hankel matrix can be represented as a sum of rank-one matrices, and linear independence of exponentials implies that its rank is equal to *k*.

Let the time series *g*(*t*) = (*f*_1_, *f*_2_, …, *f*_*N*_) be generated by Equation 6.

Consider the piecewise linear trajectory *K*_*g*(*t*)_ with embedding window *n*, where *k* ≤ *n* ≤ *N* − *n* + 1:

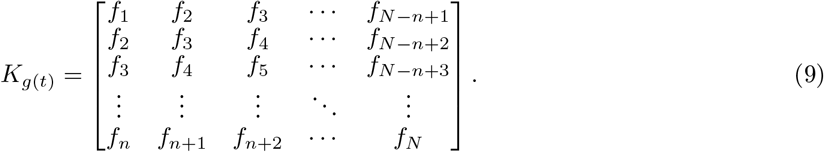

The columns of this matrix are vectors in ℝ^*n*^:

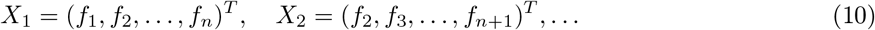

Using the representation of *g*(*t*), each column can be written as

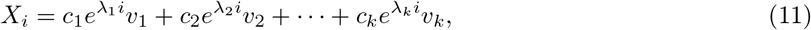

where

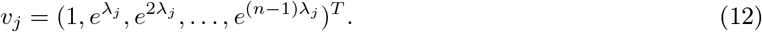

Thus, all points of the trajectory *K*_*g*(*t*)_ lie in the *k*-dimensional subspace *L* spanned by {*v*_1_, …, *v*_*k*_}:

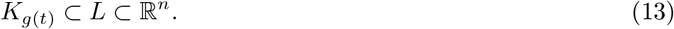

The dimension *k* does not exceed the rank of the corresponding Hankel matrix, which completes the proof.□

##### Corollary 1.

If prior information indicates that the time series *g*(*t*) of length *N* is generated by at most *k* sources of the form 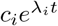, then the number of components (sources) can be determined by computing the rank of the Hankel matrix *H*_*g*(*t*)_ with embedding window *n* satisfying *k* ≤ *n* ≤ *N* − *n* + 1.

The dimension of the space containing the trajectory *K*_*g*(*t*)_ therefore directly reflects the number of underlying exponential components generating the signal.

### Sum of harmonic components

Within the geometric framework of [7, 8, 9, 10], we characterize the dimension of the subspace in which the piecewise linear trajectory of a time series lies when the signal is generated by a superposition of *k* harmonic components of the form Equation 14.

We now analyze the geometry of delay embeddings for time series generated by sums of harmonic components of the form

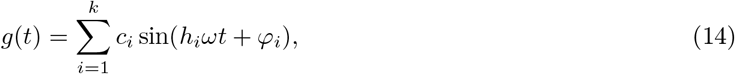

where *c*_*i*_, *ω, h*_*i*_, and *φ*_*i*_ are constant parameters.

#### Lemma 2

(Subspace structure of harmonic signals). The delay embedding of a time series *g*(*t*) defined by Equation 14 lies in a 2*k*-dimensional subspace

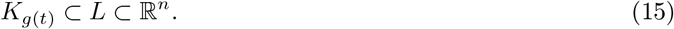

Moreover, this subspace is spanned by pairs of vectors

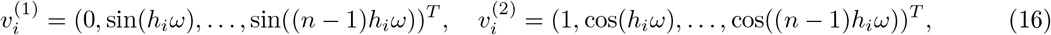

for *i* = 1, …, *k*.

Each harmonic component defines a two-dimensional plane span 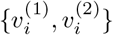, and the embedding is contained in the direct sum of these planes.

Proof▷ Using the trigonometric identity

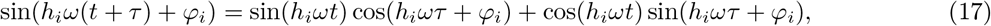

each shifted signal can be expressed as a linear combination of sin(*h*_*i*_*ωt*) and cos(*h*_*i*_*ωt*).

Thus, each column of the Hankel embedding can be written as a linear combination of vectors 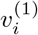 and 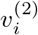. Since there are *k* independent harmonic components, the embedding lies in a subspace of dimension at most 2*k*.□

### Geometric interpretation

Each harmonic component corresponds to a two-dimensional invariant plane in the embedding space. The full trajectory is contained in the direct sum of these planes, yielding a 2*k*-dimensional structure.

The relative orientation of these planes is determined by the frequencies *h*_*i*_*ω* and can be characterized by the angles between their normal vectors.

#### Corollary 2.

If the time series *g*(*t*) is generated by at most *k* independent harmonic components, then the number of components can be recovered from the rank *r* of the Hankel embedding as

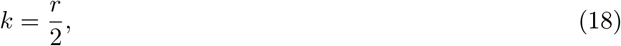

provided that the embedding window satisfies 2*k* ≤ *n* ≤ *N* − *n* + 1.

### Projection properties

The projection of the embedding onto each plane span 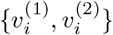 traces a closed curve. In the ideal case, this curve is inscribed in a circle.

When the sampling period is commensurate with the signal period, i.e., when

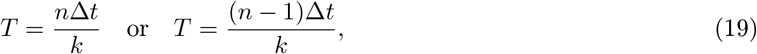

the projection forms regular polygonal structures in the corresponding plane (Fig. 4).

### Exponentially modulated oscillations

We now consider the general class of signals combining exponential and oscillatory components, which includes exponentially modulated oscillations.

#### Lemma 3

(Subspace structure of exponentially modulated oscillations). The delay embedding of a time series *g*(*t*) generated by

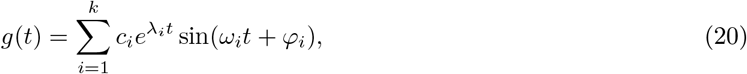

lies in a 2*k*-dimensional subspace *L* ⊂ ℝ^*n*^, i.e. *K*_*g*(*t*)_ ⊂ *L*.

Then the rank of its Hankel embedding satisfies

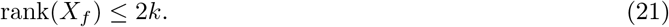

Under generic conditions on (*λ*_*i*_, *ω*_*i*_), this bound is tight.

Proof▷ Each component 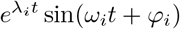 can be expressed as a linear combination of complex exponentials:

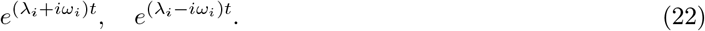

Thus, each component contributes two linearly independent directions to the embedding space.

The Hankel embedding of *g*(*t*) can be represented as a linear combination of two exponential sequences with complex conjugate exponents. Therefore, each component contributes at most two dimensions to the column space of *X*_*f*_ . The result follows by linearity. Under generic conditions on the parameters, this bound is tight, and therefore 2*k*.□

### Geometric interpretation

Each exponentially modulated oscillatory component defines a two-dimensional subspace in the embedding space, corresponding to a pair of conjugate exponential modes. The full trajectory is contained in the direct sum of these subspaces.

Thus, the geometry of the embedding combines exponential growth or decay with rotational structure, forming a family of invariant two-dimensional planes with non-uniform scaling.

### Relation to special cases

The result generalizes previous cases:

- when *ω*_*i*_ = 0, the model reduces to exponential signals with dimension *k*,
- when *λ*_*i*_ = 0, it reduces to harmonic signals with dimension 2*k*.

#### Corollary 3.

If a time series is generated by at most *k* exponentially modulated oscillatory components, then the number of components can be estimated from the rank *r* of the Hankel embedding as

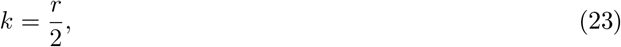

provided that the embedding window satisfies 2*k* ≤ *n* ≤ *N* − *n* + 1.

### Interpretation

This lemma establishes a direct link between the number of underlying dynamical components and the intrinsic dimension of the embedding.

This result unifies exponential and harmonic models within a single geometric framework. The intrinsic dimension of the embedding reflects the number of oscillatory components, while the structure of the corresponding subspaces encodes both frequency and growth/decay characteristics of the signal.

#### Theorem 1

(Geometric invariance of structured time series embeddings). Let a time series *g*(*t*) be generated by a finite superposition of dynamical components of the form

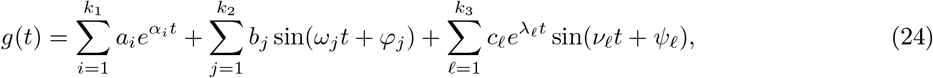

where all parameters are real and all components are pairwise non-degenerate.

Let *H*_*g*(*t*)_ denote the Hankel embedding of *g*(*t*) with embedding window *n*, and let *K*_*g*(*t*)_ be the associated piecewise linear trajectory.

Then there exists a linear subspace *L* ⊂ R^*n*^ such that

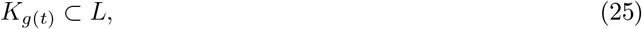

and the dimension of *L* satisfies

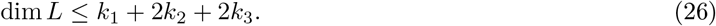

Moreover, this dimension is invariant with respect to sampling, embedding, and observation noise (up to *ε*-perturbations of the subspace), and uniquely characterizes the generative structure of the signal.

Proof▷ The result follows by superposition of the three cases established above.

Each pure exponential term 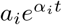 contributes one independent direction to the embedding space. Each harmonic term *b*_*j*_ sin(*ω*_*j*_*t* + *ϕ*_*j*_) contributes a two-dimensional invariant plane generated by the corresponding sine and cosine modes. Each exponentially modulated oscillatory term 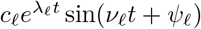 contributes two independent directions corresponding to the conjugate exponential modes 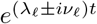.

Since the Hankel embedding is linear with respect to the signal, the embedding of *g*(*t*) is contained in the sum of the corresponding subspaces. Therefore, the dimension of the ambient subspace does not exceed

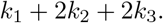

Under generic conditions, these directions are linearly independent, so equality holds.□

### Interpretation

The theorem establishes a direct correspondence between the generative mechanism of a time series and the geometry of its delay embedding. Each exponential component contributes one dimension, while each oscillatory component contributes a two-dimensional invariant plane, reflecting the presence of conjugate dynamical modes.

As a consequence, the intrinsic dimension of the embedding is not merely a statistical property but an invariant of the underlying dynamics. This result implies that time series generated by fundamentally different mechanisms occupy distinct regions of the Grassmann manifold, even when their spectral or statistical characteristics are indistinguishable.

This correspondence provides a complete geometric representation of a broad class of dynamical systems and forms the theoretical foundation for classification via subspace separation.

### Implication

This theorem provides a complete geometric characterization of a broad class of dynamical systems. In particular, it establishes that the intrinsic dimension of delay embeddings is an invariant of the underlying generative mechanism, rather than a property of the observed data.

This result elevates delay embedding from a reconstruction tool to a representation in which dynamics become geometrically observable.

### Effect of the embedding window

The sensitivity of the proposed method can be controlled through the choice of the embedding window *n*.

An important characteristic of the trajectory *K*_*g*(*t*)_ for signals of the form Equation 6 that depends on *n* is the cosine of the angle between vectors

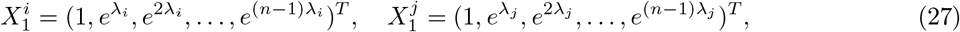

corresponding to components of a signal of the form (Equation 6). This cosine is given by

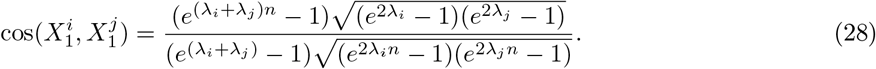

Consider the case *λ*_*i*_ = −*λ*_*j*_ as *n* → ∞. Then

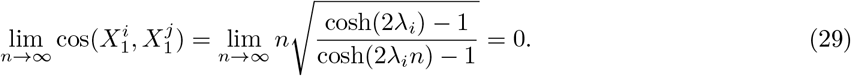

Thus, as the embedding window *n* increases, the vectors 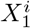 and 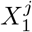 with *λ*_*i*_ = −*λ*_*j*_ rapidly become orthogonal.

### Implication

This effect enhances the separability of components in the embedding space: increasing *n* leads to improved geometric discrimination between different exponential modes. This behavior is illustrated in simulation results (see Fig. 3).

We note that for signals of the form Equations 14 and 20, it is sufficient to choose the embedding window

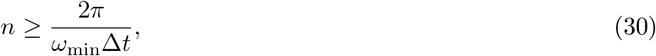

in order to recover the characteristic circular projections associated with harmonic components, where *ω*_min_ denotes the minimal frequency. See Fig. 9.

As follows from Lemma 2, a signal generated by a superposition of independent harmonic components Equation 14, is characterized by the fact that the projection of the embedding *K*_*g*(*t*)_ onto the plane associated with the *i*-th harmonic coincides with the projection of the embedding of the individual harmonic component. In particular, this projection is inscribed in a circle (Fig. 7).

At the same time, the embedding of each individual harmonic lies in the corresponding two-dimensional subspace spanned by the vectors 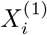 and 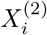, for all *n, k, N* such that 2*k* ≤ *n < N* .

In the model case where the Fourier spectrum is discrete and consists of frequencies that are integer multiples of a minimal frequency *ω*_min_, the characteristic geometric structure can be recovered by choosing the embedding window

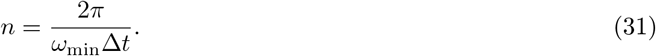

If the signal contains a harmonic with period exceeds the embedding window defined by Equations 14, the projections lose their regular polygonal structure (Fig. 8).

### Topological classifier

The proposed approach treats time series classification as a problem of geometric structure identification. A time series is mapped to a trajectory in a high-dimensional space via delay embedding, and its class is determined by the low-dimensional subspace in which this trajectory resides.

Thus, classification is reduced to identifying the geometric form of the embedding rather than analyzing signal values or spectral features.

### Grassmannian representation

Let *r* = rank(*X*_*f*_). We associate each time series with a point on the Grassmann manifold:

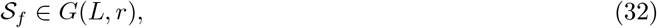

where *G*(*L, r*) denotes the set of *r*-dimensional subspaces of ℝ^*L*^.

Thus, time series classification is reformulated as a problem of classifying points on a Grassmann manifold.

### Class subspaces

Let ℱ = {*f*_1_, …, *f*_*n*_} be a collection of time series with class labels *y*_*i*_ ∈ {1, …, *M*}. For each class *c*, we construct a representative subspace *L*_*c*_ ∈ *G*(*L, r*) by solving the optimization problem:

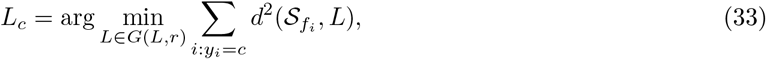

where *d*(·, ·) is a Grassmannian distance (e.g., projection or geodesic distance).

To enhance separation between classes, we introduce a max–min criterion:

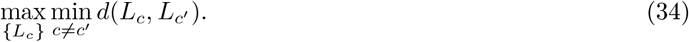

Topological classifier Principals and Algorithms

For each class *c*, define an *ε*_*c*_-neighborhood:

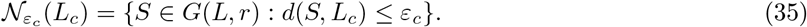

#### Definition 1

(Topological classifier). Given a time series *f*, its class label is defined as

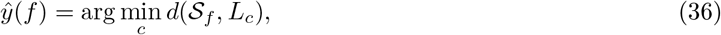

subject to

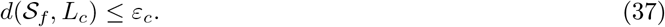

If no class satisfies the tolerance condition, the sample may be rejected or assigned to the closest class.

#### Separation property

The classifier relies on the assumption that embeddings of different classes concentrate near distinct subspaces.

##### Proposition 1

(Subspace separability). Assume that for each class *c*, all embeddings satisfy

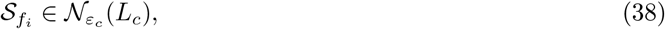

and that for any *c* ≠ *c*^*′*^,

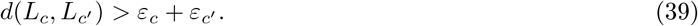

Then the classifier *ŷ*(*f*) is well-defined and yields zero classification error on the training set.

Proof▷ The separation condition ensures that neighborhoods do not intersect. Therefore, each embedding belongs to exactly one class neighborhood, and the nearest-subspace rule recovers the correct label.□

### Interpretation

The proposed classifier assigns labels based on geometric proximity in the embedding space. Thus, classification depends on the topology of the reconstructed trajectories rather than on pointwise signal features or spectral coefficients.

### Problem formulation

Let ℱ = {*f*_1_, …, *f*_*k*_} be a collection of time series. Each *f*_*i*_ is mapped to an embedded representation *X*_*i*_ ⊂ ℝ^*n*^. The goal is to construct a family of subspaces {*L*_1_, …, *L*_*m*_} of dimension *r* and corresponding tolerances {*ε*_1_, …, *ε*_*m*_} such that:

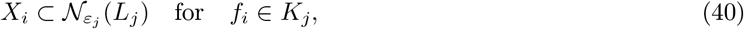

where 𝒩_*ε*_(*L*) denotes the *ε*-neighborhood of subspace *L*.

#### Algorithm 1

Formation of Equivalence Classes of Time Series via Topological Separation

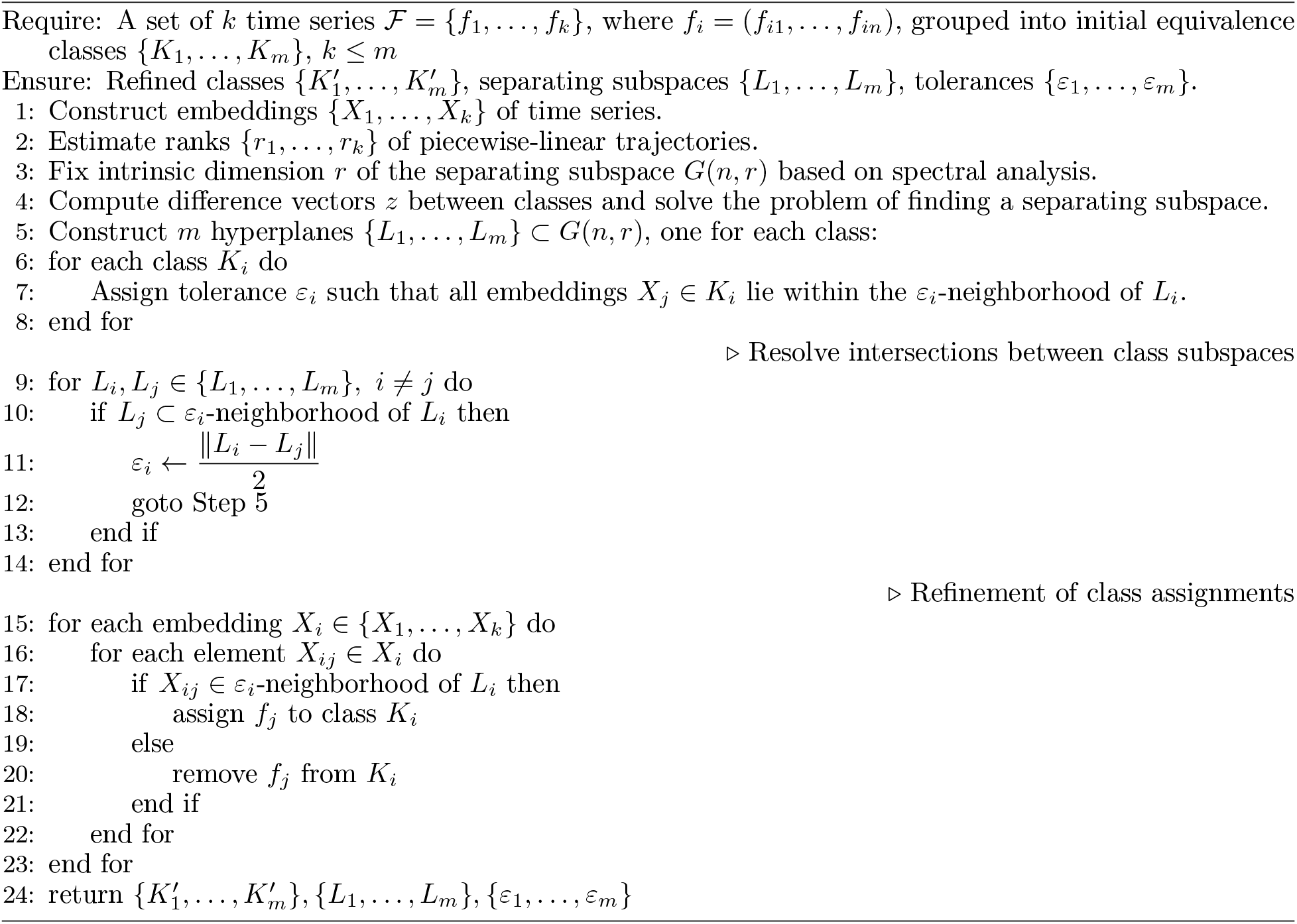

#### Algorithm 2

Classification in embedding space

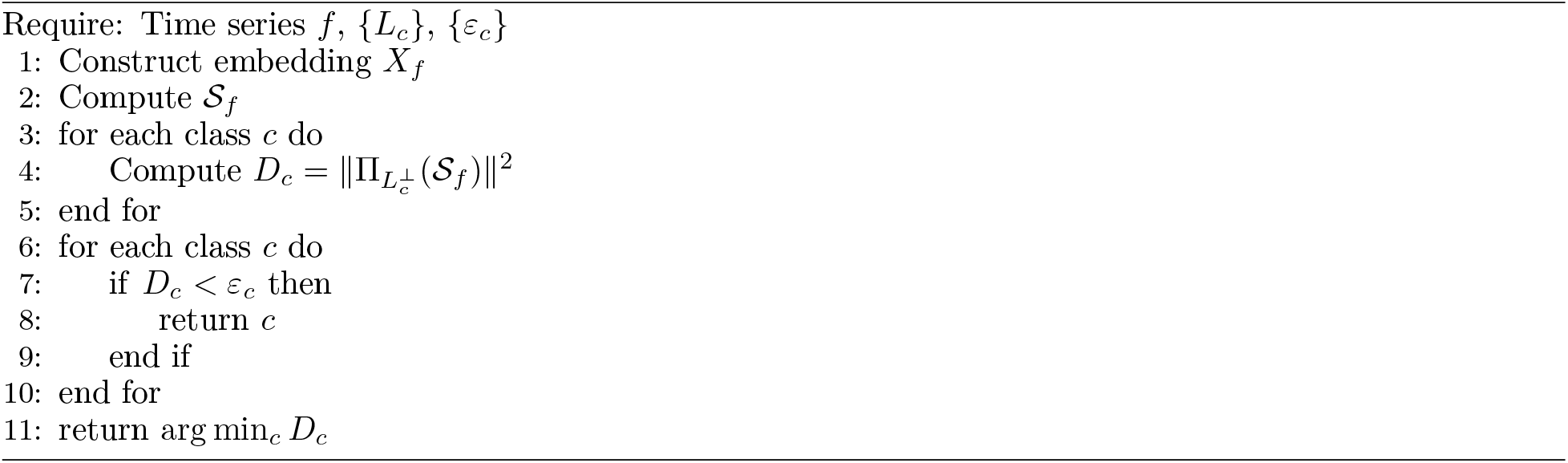

#### Summary

The proposed method combines:

- low-rank structure of Hankel embeddings,
- Grassmannian geometry,
- and subspace-based classification.

This yields a classifier that is both interpretable and grounded in the dynamics of the underlying signal.

The computational cost is dominated by the SVD of the Hankel matrix, resulting in complexity *O*(*n*^2^*N*). Distance computations on the Grassmann manifold scale with the intrinsic dimension of the subspaces.

### Effect of noise

The presence of noise can be interpreted geometrically as a perturbation of the delay embedding. Let *H*_*f*_ be the Hankel embedding of a signal and 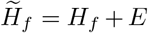 its noisy counterpart. While noise perturbs individual entries of the embedding, it does not fundamentally alter its low-dimensional structure for moderate perturbations.

As a result, the piecewise linear trajectory of the noisy signal does not leave the underlying subspace but instead forms an *ε*-neighborhood around the trajectory of the clean signal. In this sense, noise induces an *ε*-neighborhood of the underlying subspace in the embedding space centered around the original low-dimensional subspace.

This observation justifies the use of *ε*-neighborhoods in the proposed classifier: signals generated by the same dynamical mechanism are expected to occupy a common subspace together with its noise-induced neighborhood.

#### Proposition 2

(Stability of subspace under noise). Let *H*_*f*_ be the Hankel embedding of a signal *f*, and let 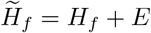 be its noisy counterpart, where *E* is a perturbation matrix with sufficiently small norm. Let 𝒮_*f*_ and 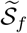 denote the corresponding column subspaces.

Then the distance between 𝒮_*f*_ and 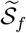 on the Grassmann manifold satisfies

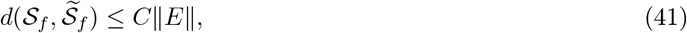

for some constant *C* depending on the spectral gap of *H*_*f*_ .

In particular, the trajectory of the noisy signal remains within an *ε*-neighborhood of the original subspace, where *ε* = 𝒪 (∥*E*∥).

Proof▷ The result follows from standard perturbation theory for invariant subspaces. Small perturbations of the matrix *H*_*f*_ lead to small changes in its principal subspace, provided that the singular values exhibit a non-degenerate gap.

Thus, the subspace 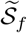 remains close to 𝒮_*f*_ in the Grassmannian distance, which implies that the corresponding trajectory lies within an *ε*-neighborhood of the original subspace.□

This result provides a theoretical justification for the use of *ε*-neighborhoods in the proposed classifier. Evaluation on synthetic and real-world data

We evaluate the proposed topological classifier on both synthetic and real-world time series in order to assess its ability to distinguish signals generated by different dynamical mechanisms.

### Synthetic data

Synthetic signals were constructed with randomly sampled parameters and additive Gaussian noise. We considered three classes of signals:

- Exponential signals (Eq. 6),
- Harmonic signals (Eq. 14),
- Exponentially modulated oscillatory signals (Eq. 20).

To assess the discriminative power of the proposed approach, we compared it with Fourier-based representations. In multiple instances, signals with distinct generative mechanisms exhibited similar spectral profiles, rendering them indistinguishable in the frequency domain. In contrast, their delay embeddings occupied distinct low-dimensional subspaces, enabling correct classification.

For each class, we estimated representative subspaces *S*_*c*_ and computed pairwise distances on the Grassmann manifold. We observed consistent separation between classes, satisfying the condition (Eq. 2), which justifies the use of subspace-based classification.

To evaluate robustness, we introduced additive Gaussian noise with increasing variance. The classifier retained correct predictions as long as the perturbations did not significantly distort the underlying subspace structure, in agreement with theoretical stability results.

### Real-world data

We further evaluated the method on a publicly available EEG dataset from the UCI Machine Learning Repository [19]. The dataset consists of 64-channel EEG recordings from 122 subjects divided into two groups: individuals diagnosed with chronic alcoholism and a control group.

For each subject, 120 trials of duration 1 second were recorded under visual stimulus conditions (Snodgrass and Vanderwart image set). Signals were sampled at 256 Hz, resulting in time series of length 256 per channel. We compared the performance of the proposed method with previously reported results on the same dataset. All experiments were repeated across multiple random realizations to ensure stability of the results.

### Experimental protocol

Data splitting and evaluation procedure. For both synthetic and real-world datasets, we employed a stratified evaluation protocol to ensure reproducibility and robustness of the results [20].

For synthetic data, signals were generated independently for each experiment using randomly sampled parameters. The dataset was split into training and testing subsets in an 80/20 ratio. To reduce variance, all experiments were repeated over multiple independent realizations (typically 10 runs), and the reported results correspond to the mean performance.

For real-world EEG data, we adopted a subject-wise evaluation protocol to avoid data leakage. Specifically, subjects were partitioned into training (80%) and testing (20%) groups, ensuring that no trials from the same subject appeared in both sets. This setup reflects a realistic scenario of generalization to unseen individuals.

Preprocessing. All time series were standardized prior to embedding by removing the mean and normalizing the variance. No additional feature engineering or filtering was applied, in order to evaluate the method in a minimally processed setting.

Embedding and parameter selection. The embedding window *n* was selected based on the minimal characteristic scale of the signal, following the condition (Eq. 30), where applicable. In practice, multiple values of *n* were evaluated, and stable performance was observed within a broad range.

Subspace estimation. For each class, representative subspaces *L*_*c*_ were estimated from the training data using singular value decomposition of the corresponding Hankel embeddings. The intrinsic dimension of each subspace was determined via rank estimation, consistent with the theoretical results.

Classification rule. Given a test signal, its delay embedding was constructed and projected onto candidate subspaces. Classification was performed by assigning the signal to the class which subspace minimized the Grassmannian distance, subject to the separability condition (Eq. 39).

Evaluation metrics. Classification performance was measured using accuracy:

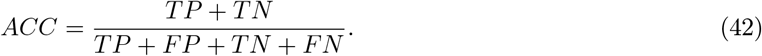

In addition, variability across runs was assessed using standard deviation.

Baseline comparison. To assess the advantages of the proposed approach, we compared it with Fourier-based representations and previously reported results on the same dataset. All comparisons were performed under comparable experimental conditions.

### Summary of the method

The proposed framework maps time series to subspaces in embedding space and performs classification via geometric separation. This replaces feature-based representations with intrinsic geometric descriptors determined by the underlying dynamics.

## Results

### Geometric structure of delay embeddings

We begin by examining the geometric structure of delay embeddings for signals with controlled generative mechanisms. Across all considered cases, we observe a consistent phenomenon: the embedding of a time series concentrates near a low-dimensional linear subspace which dimension reflects the number of underlying dynamical components.

For purely exponential signals, embeddings collapse onto one-dimensional structures, while superpositions of exponentials span higher-dimensional subspaces with dimension equal to the number of components. In contrast, harmonic signals generate two-dimensional invariant planes, characterized by circular or polygonal projections in principal component coordinates. Exponentially modulated oscillations combine these effects, producing spiral structures that encode both oscillatory and exponential dynamics.

Crucially, these geometric structures remain stable under substantial noise, with perturbations manifesting as *ε* −neighborhoods around the underlying subspaces. This stability enables reliable recovery of the intrinsic structure even in regimes where spectral methods fail to distinguish between signals.

### Sum of exponential components

To illustrate the proposed approach, we consider the delay embeddings of the following three functions:

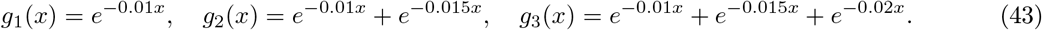

Analysis of the raw time series (Fig. 1(a)), Fourier spectra (Fig. 1(b)), and principal components does not allow one to determine whether the signal is generated by one, two, or three exponential components.

**Fig. 1:**
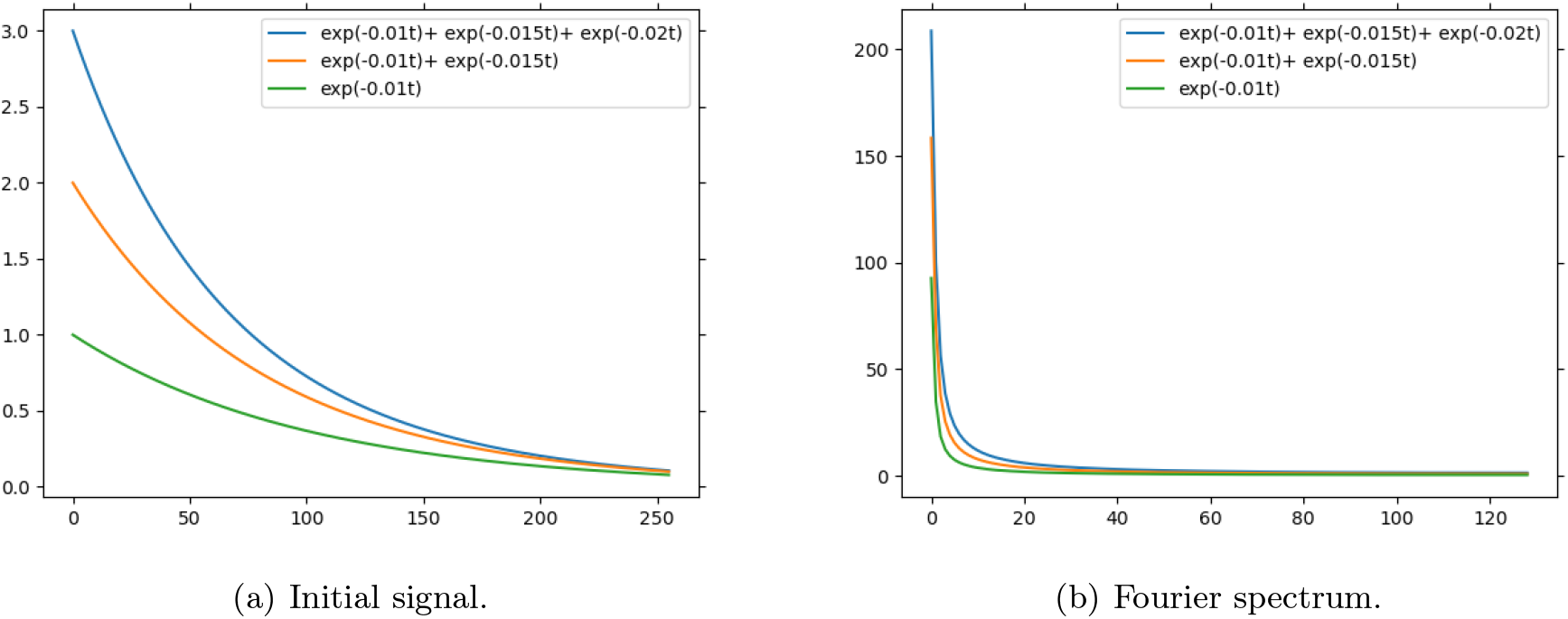
Time series corresponding to *g*_1_(*x*) = *e*^−0.01*x*^ (green), *g*_2_(*x*) = *e*^−0.01*x*^ + *e*^−0.015*x*^ (orange), and *g*_3_(*x*) = *e*^−0.01*x*^ + *e*^−0.015*x*^ + *e*^−0.02*x*^ (blue), together with their discrete Fourier spectra.

In contrast, Fig. 2(a) shows that the delay embedding of the time series generated by *g*_1_(*x*) = *e*^−0.01*x*^ lies on a one-dimensional subspace, i.e., along a line, in agreement with Lemma 1.

**Fig. 2:**
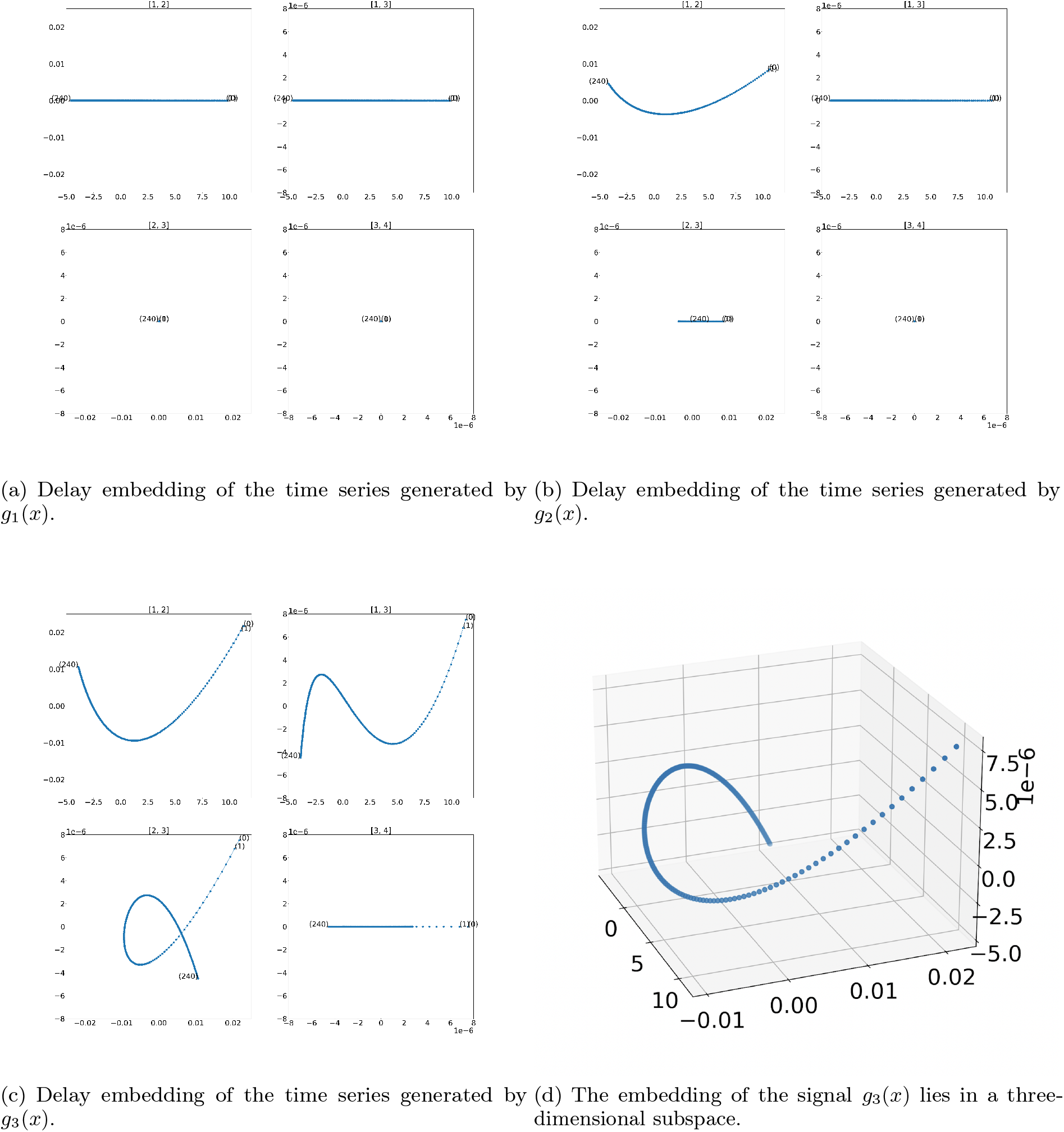
Delay embeddings of the functions *g*_1_(*x*) = *e*^−0.01*x*^, *g*_2_(*x*) = *e*^−0.01*x*^ + *e*^−0.015*x*^, and *g*_3_(*x*) = *e*^−0.01*x*^ + *e*^−0.015*x*^ + *e*^−0.02*x*^, projected onto planes spanned by pairs of principal components. Parameters: embedding dimension *n* = 16, sampling step Δ*t* = 1, initial time *t*_1_ = 0, and series length *N* = 256.

The embedding of the signal *g*_2_(*x*) = *e*^−0.01*x*^ + *e*^−0.015*x*^ lies in a two-dimensional subspace spanned by the first two principal components (Fig. 2(b)). Projections onto planes defined by the first and third, as well as the second and third principal components, remain one-dimensional.

Finally, the embedding of the signal *g*_3_(*x*) = *e*^−0.01*x*^ + *e*^−0.015*x*^ + *e*^−0.02*x*^ lies in a three-dimensional subspace, as shown in Fig. 2(c, d).

Emphasizing the strength of the proposed topological analysis of delay embeddings, we note that Fourier spectral analysis of a time series generated by a function of the form of Equation 6 does not allow one to distinguish whether the signal is composed of a sum of exponential components or a single exponential term.

Similarly, principal component analysis of such a time series does not reveal more than two dominant components for one-dimensional signals of the form of Equation 6.

In contrast, as follows from Lemma 1, the analysis of the topology of the delay embedding makes it possible to recover information about the number of exponential components of the form 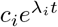 underlying the signal.

### Effect of the embedding window

The sensitivity of the proposed method can be controlled through the choice of the embedding window *n*. Fig. 3. This demonstrates how increasing the embedding window enhances the geometric separation of components.

**Fig. 3:**
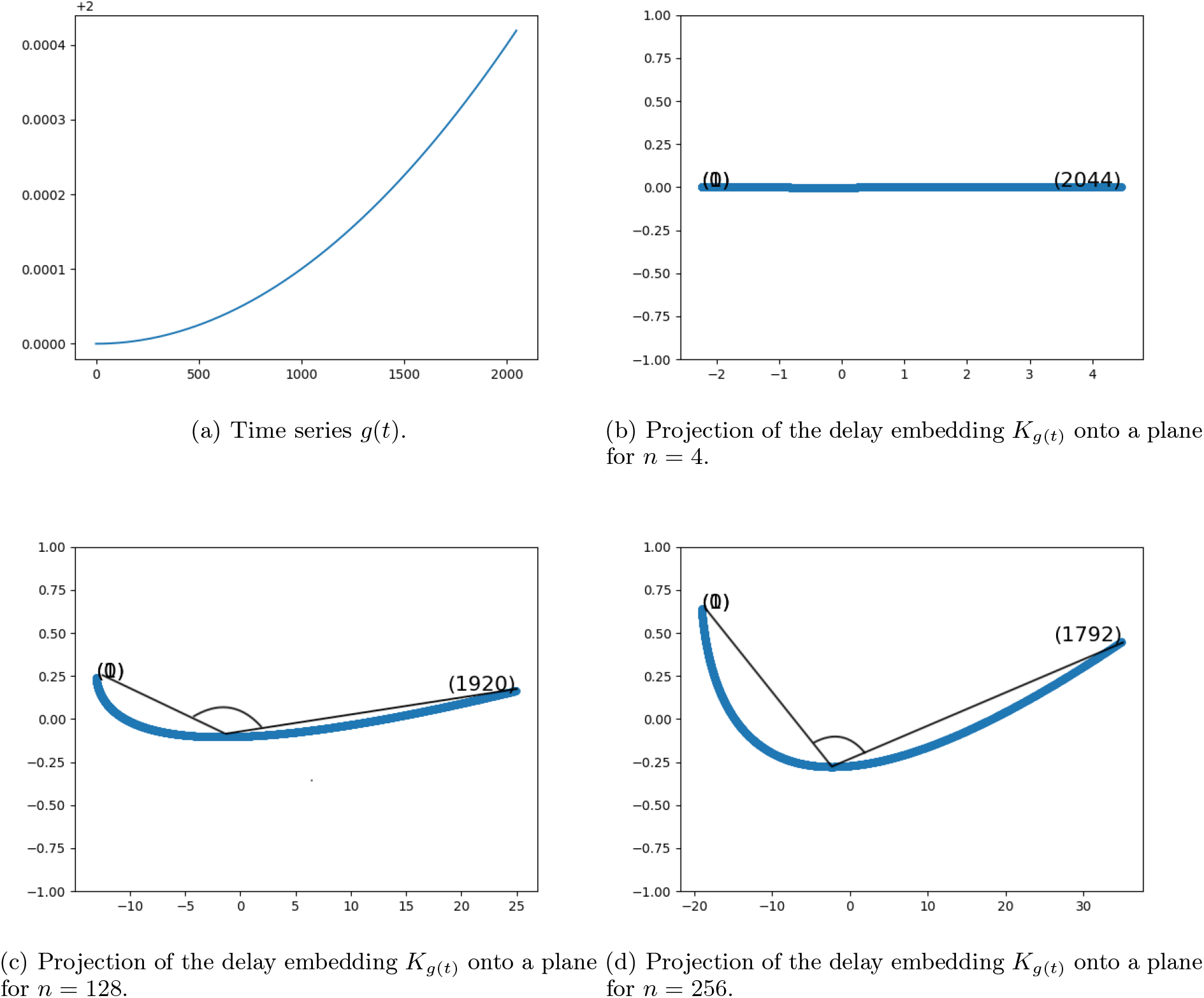
The function 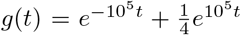 with parameters *n* = 16, Δ*t* = 1, *t*_1_ = 0, and *N* = 2048. The figure illustrates how the shape of the projection of the delay embedding *K*_*g*(*t*)_ onto the plane spanned by the first two principal components changes as the embedding window *n* varies.

### Sum of harmonic components

The projection of the delay embedding of a harmonic signal onto the plane spanned by the corresponding principal components is inscribed in a circle.

When the period of the signal satisfies Equation 19, where *k* is a divisor of *n*, the projections take the form of regular polygons with 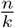 or 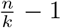 vertices, respectively, in the plane spanned by orthogonal principal components (see Fig. 4).

**Fig. 4:**
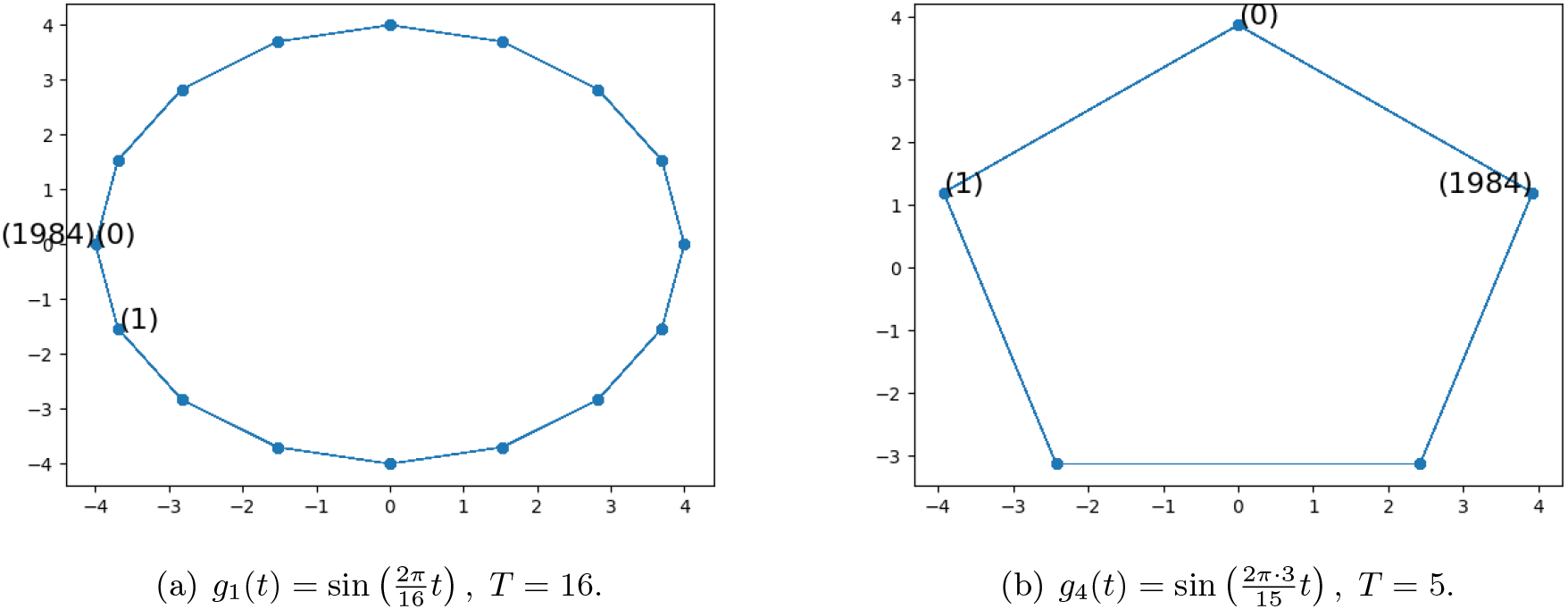
Projections of delay embeddings of the time series corresponding to 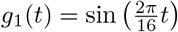 (period *T* = 16) and 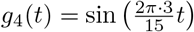 (period T = 5) onto planes spanned by pairs of principal components. The project exhibit characteristic polygonal structures, corresponding to a 16-gon and a 15-gon, respectively. Parameters: *n* = 16, Δ*t* = 1, *t*_1_ = 0, *N* = 2000.

When the parameters *k* and *n* are coprime, the *n* points of the delay embedding lie on a circle. However, the corresponding piecewise linear trajectory does not form regular polygons when projected onto planes spanned by principal components. Instead, the trajectory exhibits characteristic patterns associated with cyclic structures of order *n* and *n* − 1 generated by powers of *k* (Fig. 5(a,b)).

**Fig. 5:**
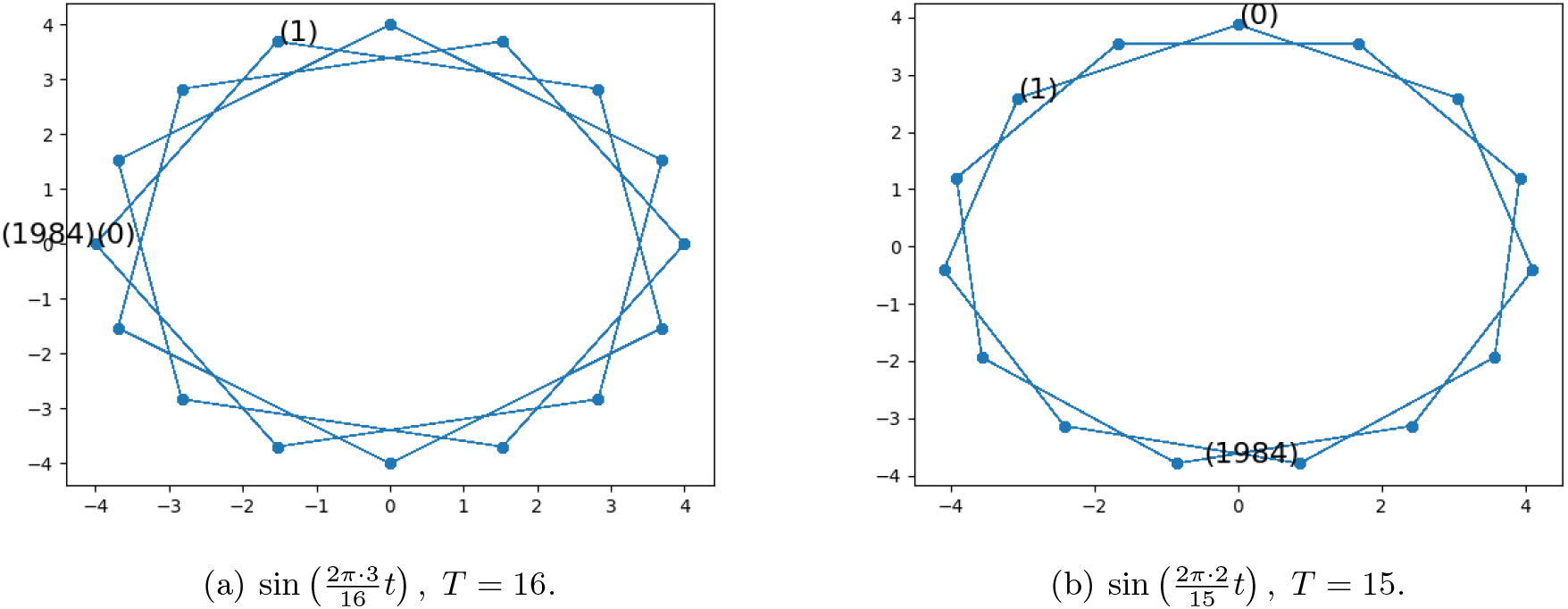
Embedding parameters: *n* = 16, Δ*t* = 1, *t*_1_ = 0, *N* = 2000.

In this regime, the trajectory may not form a closed curve. For irrational frequency ratios, the projection of the embedding converges to a circular structure as *N* → ∞ (Fig. 6). In this case, the Euclidean norm of the embedding points exhibits periodic behavior.

**Fig. 6:**
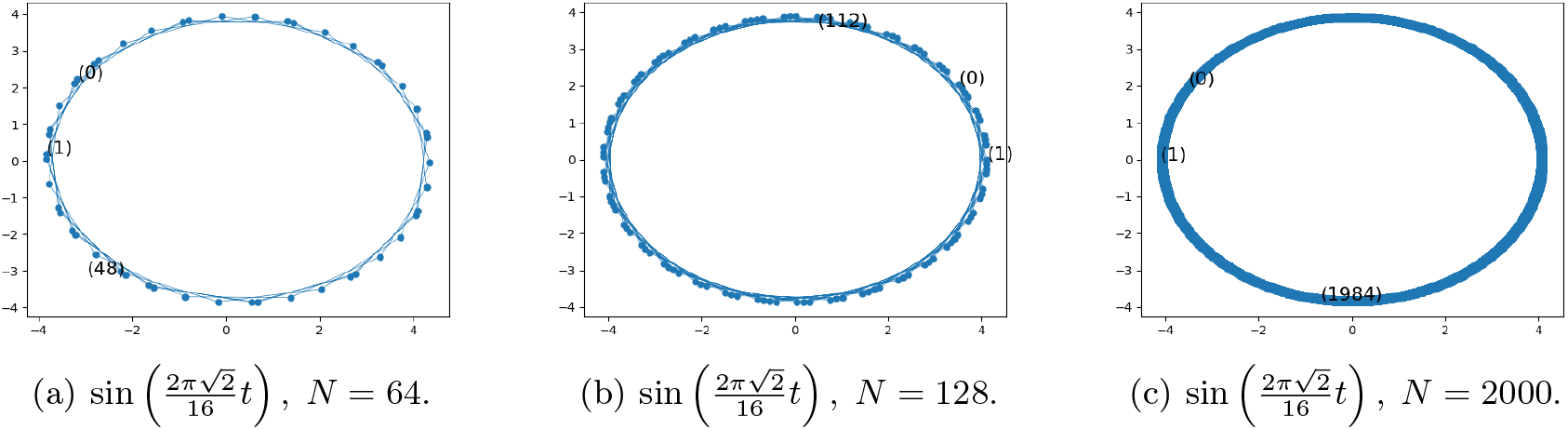
Embedding parameters: *n* = 16, Δ*t* = 1, *t*_1_ = 0.

### Effect of the embedding window

The characteristic circular projections are lost when the embedding window is below the threshold given by Equation 30. See Fig. 8. Therefore, to recover the characteristic circular projections associated with harmonic components, it is sufficient to choose the embedding window given by Equation 30. See Fig. 9.

**Fig. 7:**
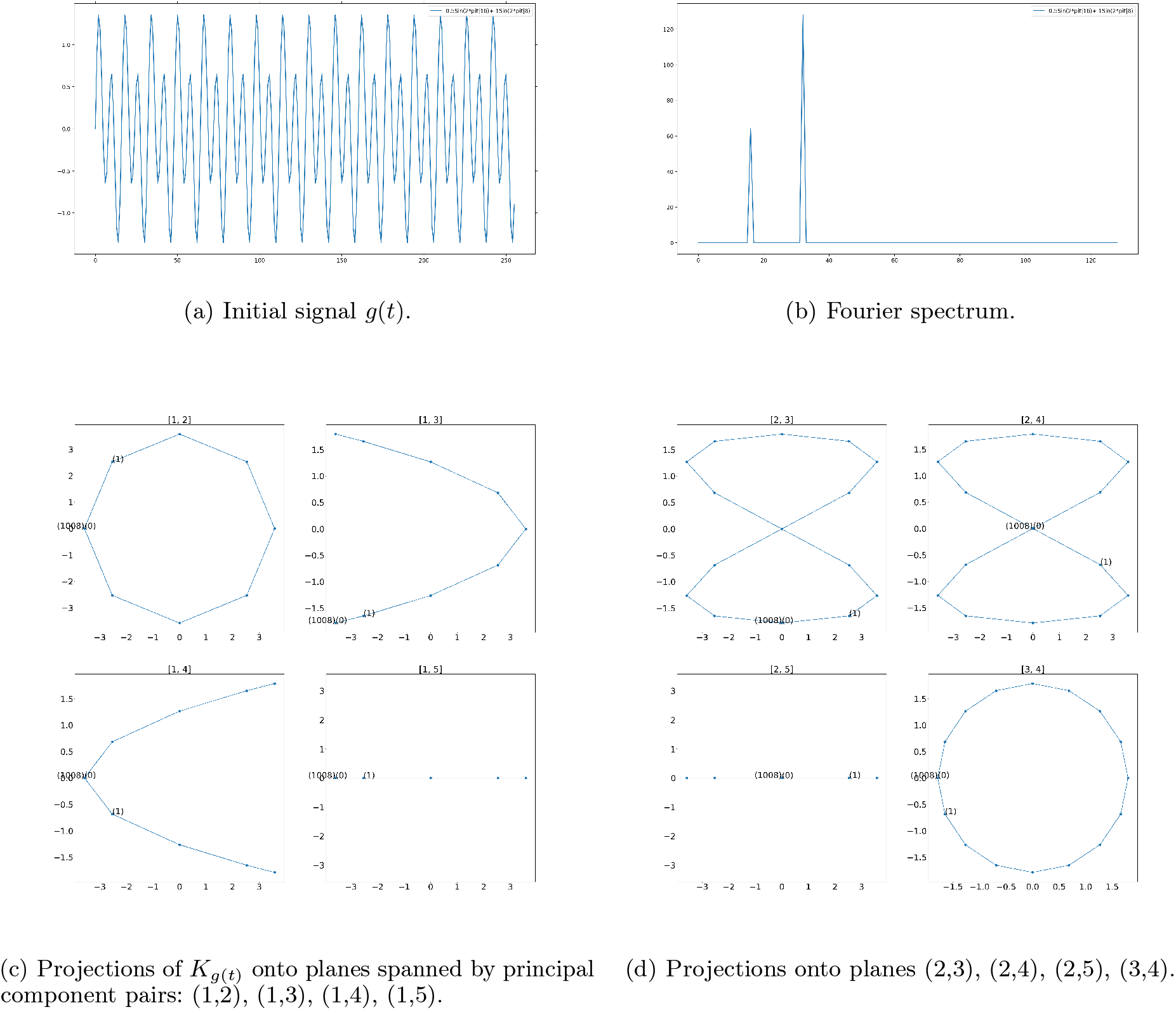
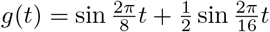, *n* = 16, Δ*t* = 1, *t*_1_ = 0, *N* = 256. Planes spanned by principal component pairs (1,2) and (3,4) correspond to the first and second harmonics and exhibit characteristic polygonal structures inscribed in circles.

**Fig. 8:**
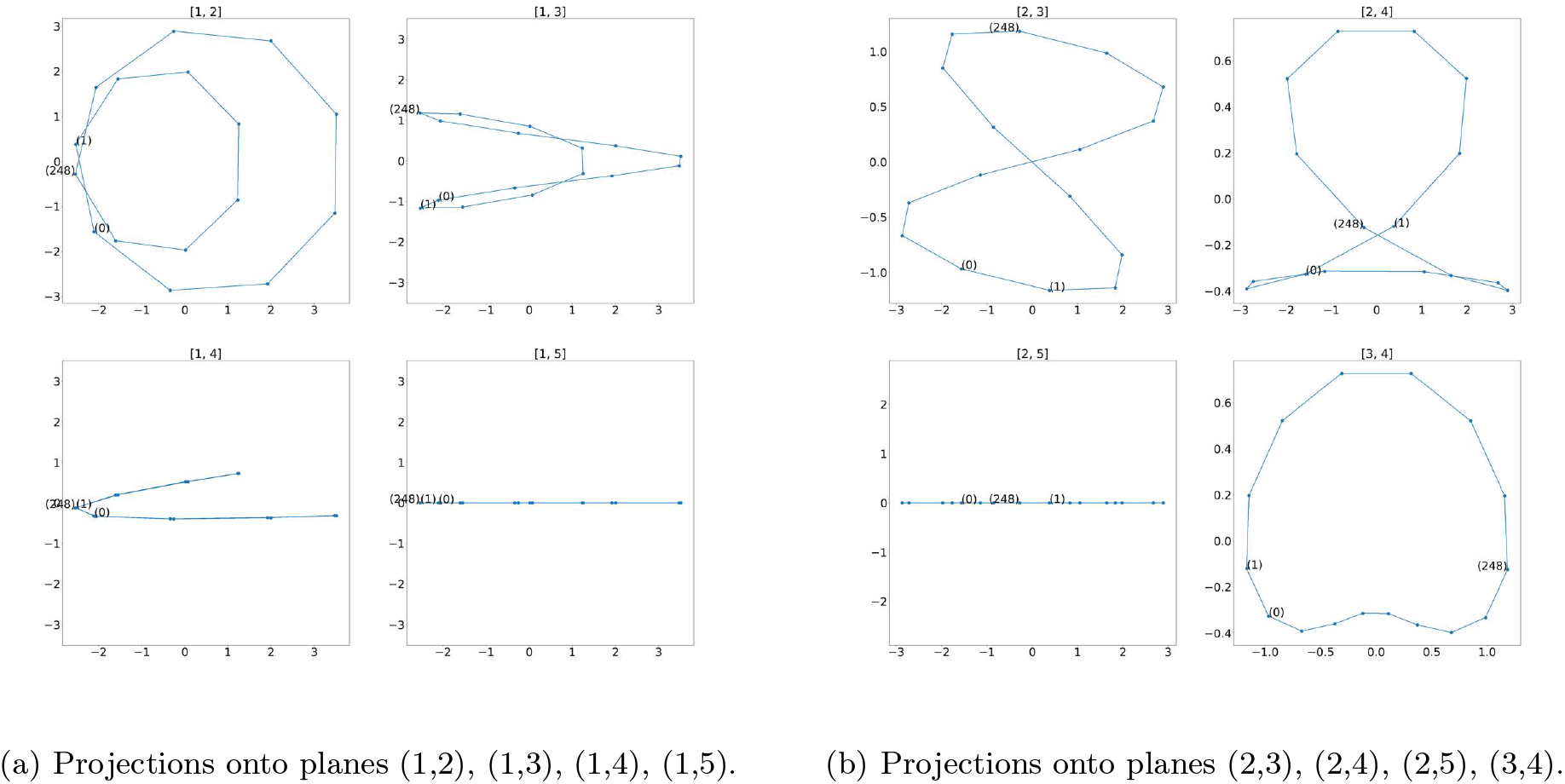
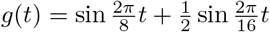, *n* = 8, Δ*t* = 1, *t*_1_ = 0, *N* = 256. The characteristic circular projections are lost when the embedding window is below 16.

**Fig. 9:**
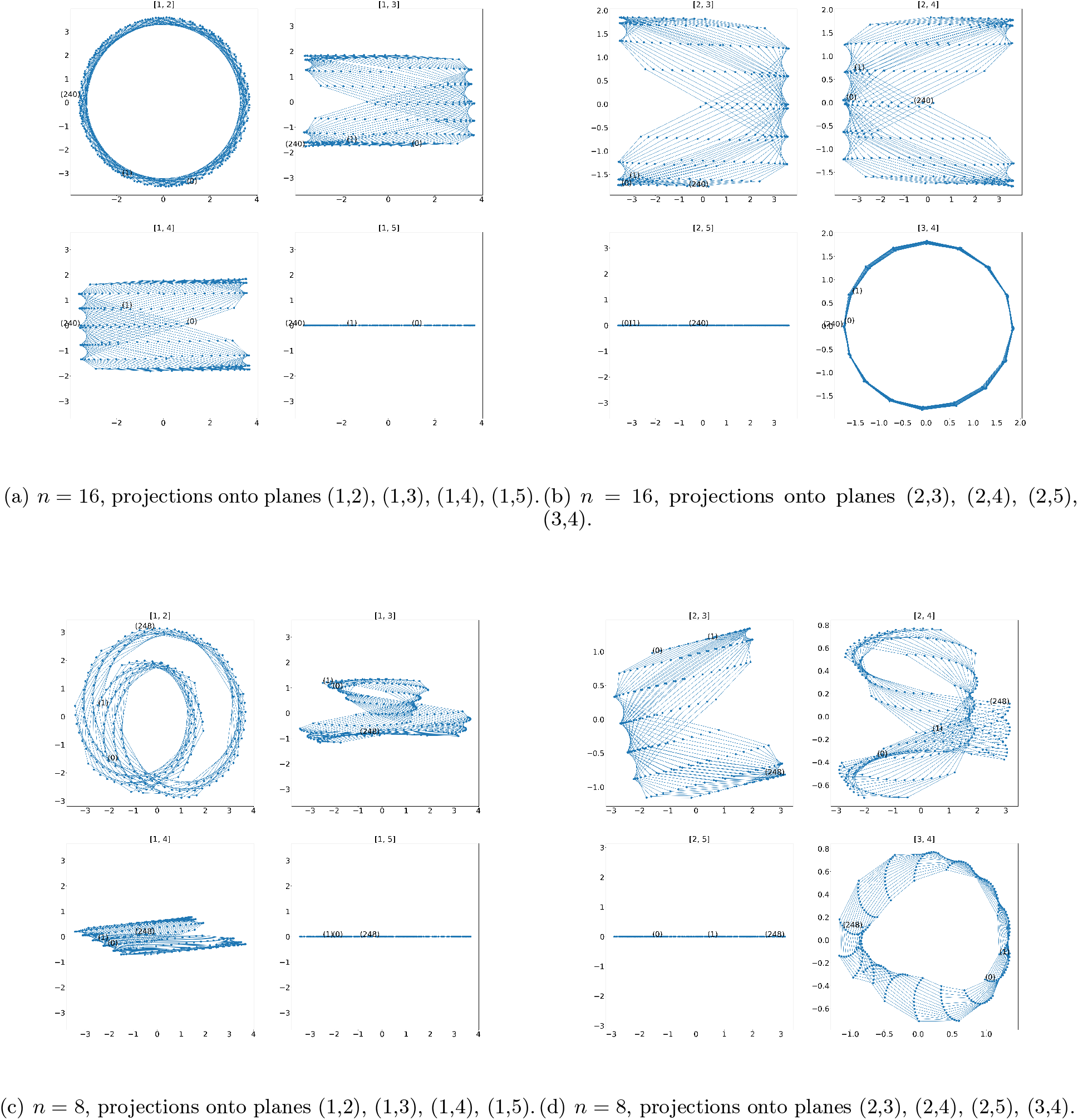
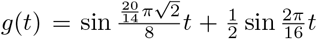, Δ*t* = 1, *t*_1_= 0, *N* = 256. Panels (a,b) show characteristic circular projections when the embedding window satisfies Equation 30; panels (c,d) show the loss of structure for smaller *n*.

### Exponentially modulated oscillations

The projection of the delay embedding of a signal approximated by a function of the form of Equation 20 onto the plane spanned by the corresponding principal components exhibits a characteristic spiral structure. In this representation, the oscillatory component determines the initial radius of the spiral, while the exponential component governs the rate of its contraction. See Fig. 10.

**Fig. 10:**
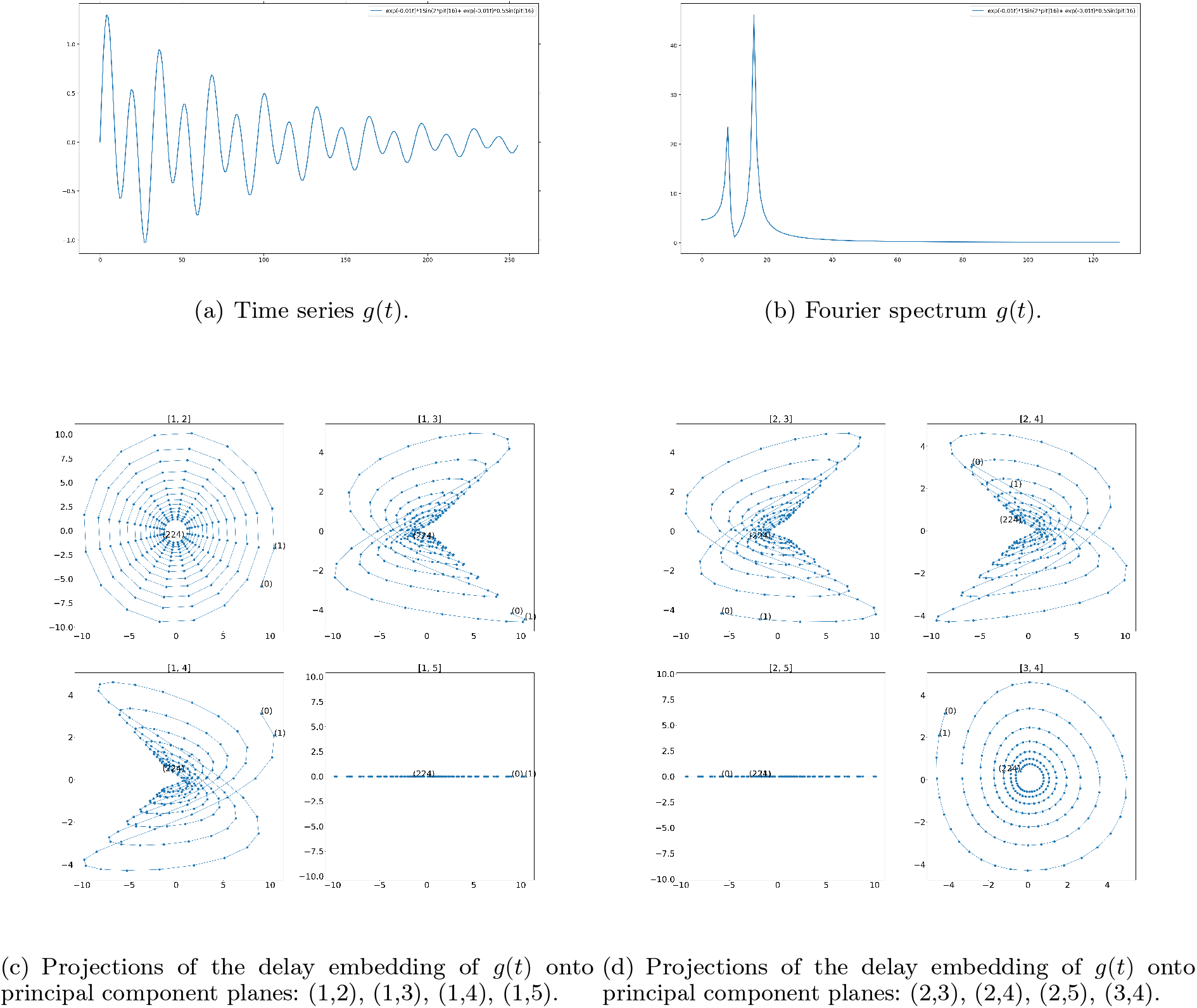
The function 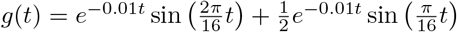, consisting of two exponentially modulated oscillatory components (with harmonic periods *T*_1_ = 16 and *T*_2_ = 8), with parameters *n* = 16, Δ*t* = 1, *t*_1_ = 0, and *N* = 256. Characteristic spiral structures associated with exponentially modulated oscillations are observed in projections onto principal component planes (1,2) and (3,4).

### Implications for classification

The observed geometric structures of delay embeddings provide a direct basis for time series classification. Signals generated by different dynamical mechanisms give rise to embeddings with distinct low-dimensional geometry, characterized by the dimension of the underlying subspace and the shape of its projections.

In particular, exponential components produce one-dimensional structures, while harmonic and exponentially modulated oscillatory components generate two-dimensional invariant planes with characteristic circular or polygonal projections. These geometric features are stable under moderate perturbations and remain identifiable even when classical methods fail to distinguish between signals.

As a result, classification can be formulated as a problem of identifying the subspace and its geometric structure in which the embedding resides. This allows one to separate signals according to their underlying generative mechanisms by comparing their positions in the space of subspaces, providing a principled and interpretable alternative to spectral and data-driven approaches.

### Effect of noise

A classical problem is the extraction of exponential/harmonic signals in the presence of Gaussian noise with known variance. We demonstrate how this problem can be addressed using the delay embedding approach. See Fig. 11.

**Fig. 11:**
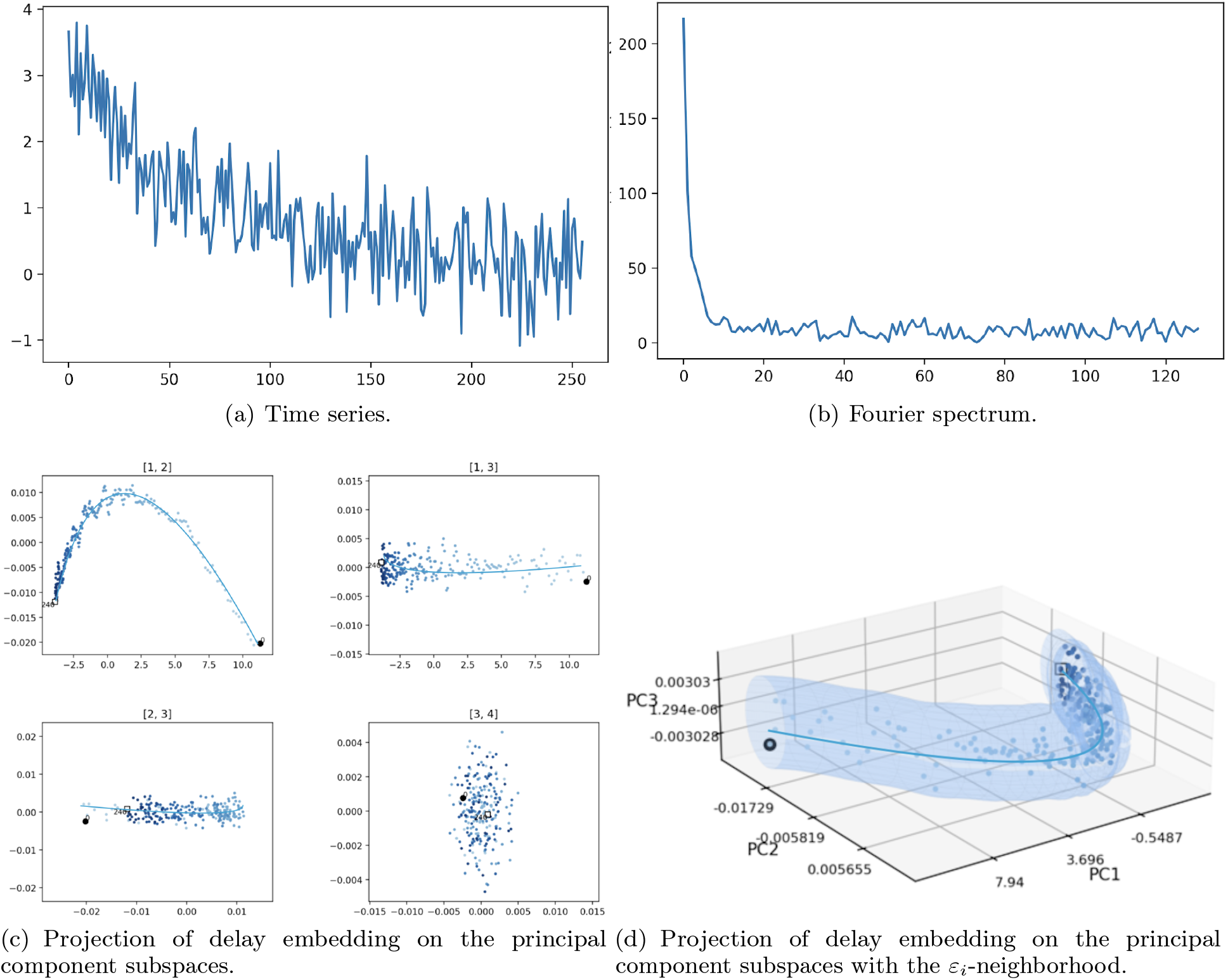
Exponential signal perturbed by additive Gaussian noise. Shown are the time series, the Fourier spectrum of eigenvalues of the scatter matrix, and projections of the delay embedding onto principal component planes (1,2), (1,3), (2,3), (3,4), for the model signal *g*(*t*) = *e*^−0.01*t*^ + *e*^−0.015*t*^ + *e*^−0.02*t*^ with Gaussian noise (*E* = 0, *D* = 0.1). Parameters: *n* = 16, Δ*t* = 1, *t*_1_ = 0, *N* = 2000.

When a harmonic signal is perturbed by additive Gaussian noise, random phase, and/or random amplitude fluctuations, the delay embedding method still allows the underlying structure of the signal to be observed in projections onto the first two principal components. In this case, the projection of the embedding retains a characteristic circular structure, albeit distorted by noise. The noise component, however, manifests itself in projections onto planes spanned by higher-order principal components. See Fig. 12.

**Fig. 12:**
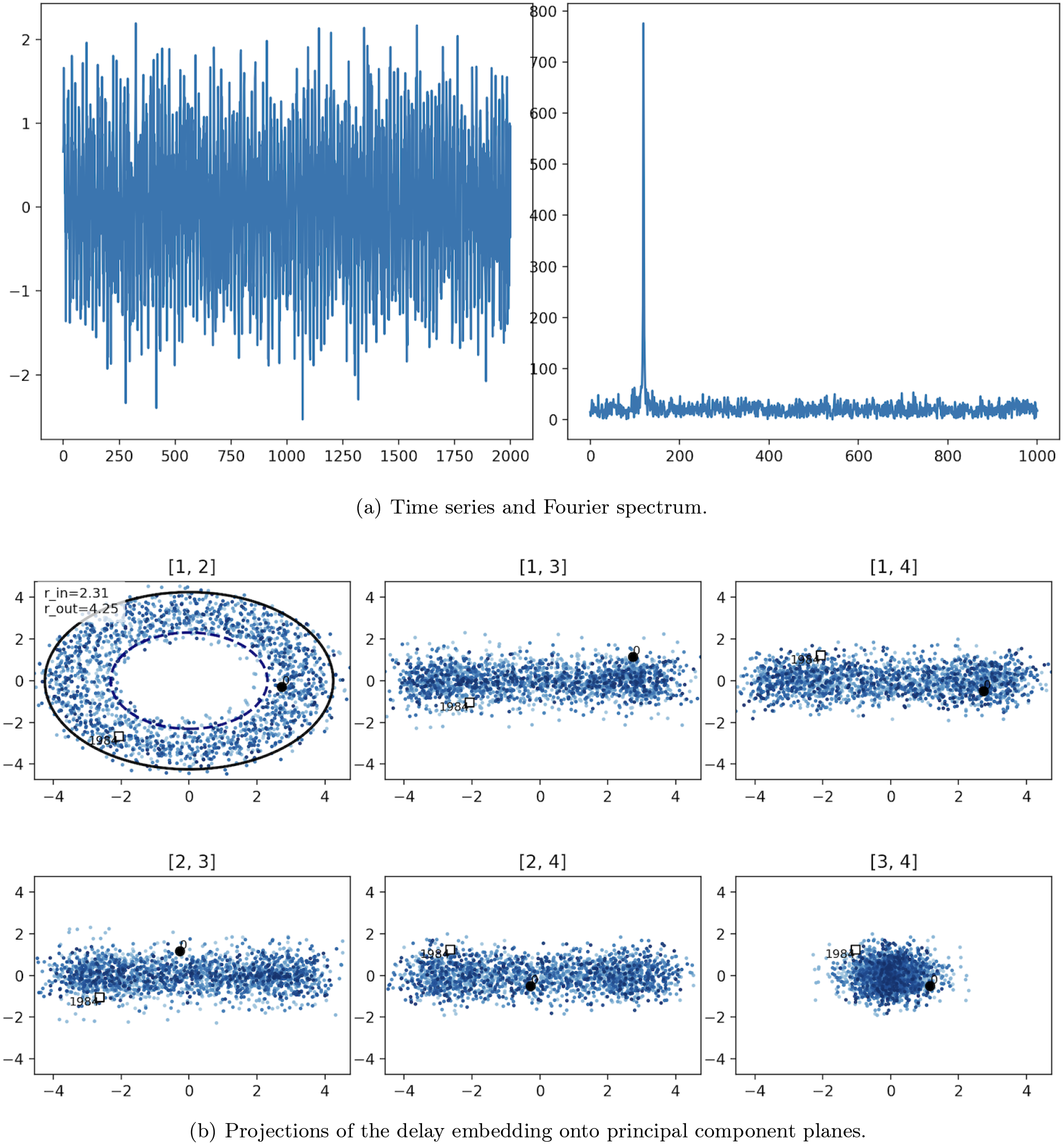
Harmonic signal perturbed by additive Gaussian noise. Shown are the time series, the spectrum of eigenvalues of the scatter matrix, and projections of the delay embedding onto principal component planes (1,2), (1,3), (2,3), (2,4), (3,4) for the model signal 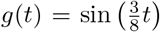 with Gaussian noise (*E* = 0, *D* = 0.5). Parameters: 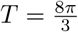, *n* = 16, Δ*t* = 1, *t*_1_ = 0, *N* = 2000.

If Gaussian noise is introduced into the argument of the harmonic signal, the resulting function becomes quasi-periodic. The delay embedding method still reveals the underlying structure in projections onto the first and second principal components. See Fig. 13.

**Fig. 13:**
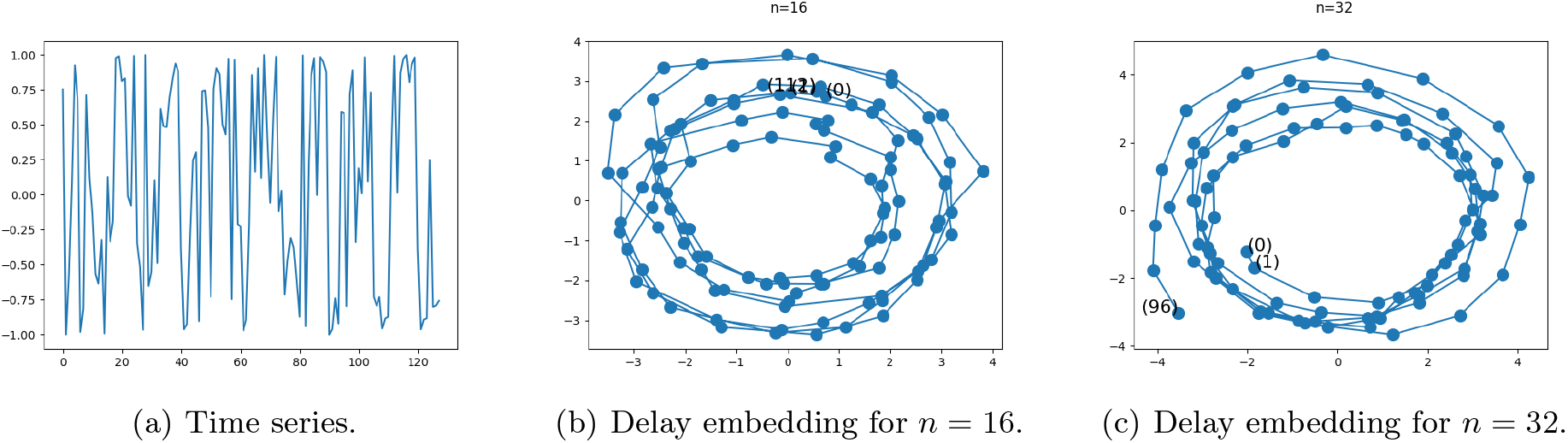
Harmonic signal with random phase perturbation: sin 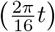, Gaussian noise (*E* = 0, *D* = 1). Parameters: Δ*t* = 1, *t*_1_ = 0, *N* = 128.

Similarly, when the signal is subject to random amplitude perturbations, the delay embedding preserves the geometric structure of the signal. See Fig. 14.

**Fig. 14:**
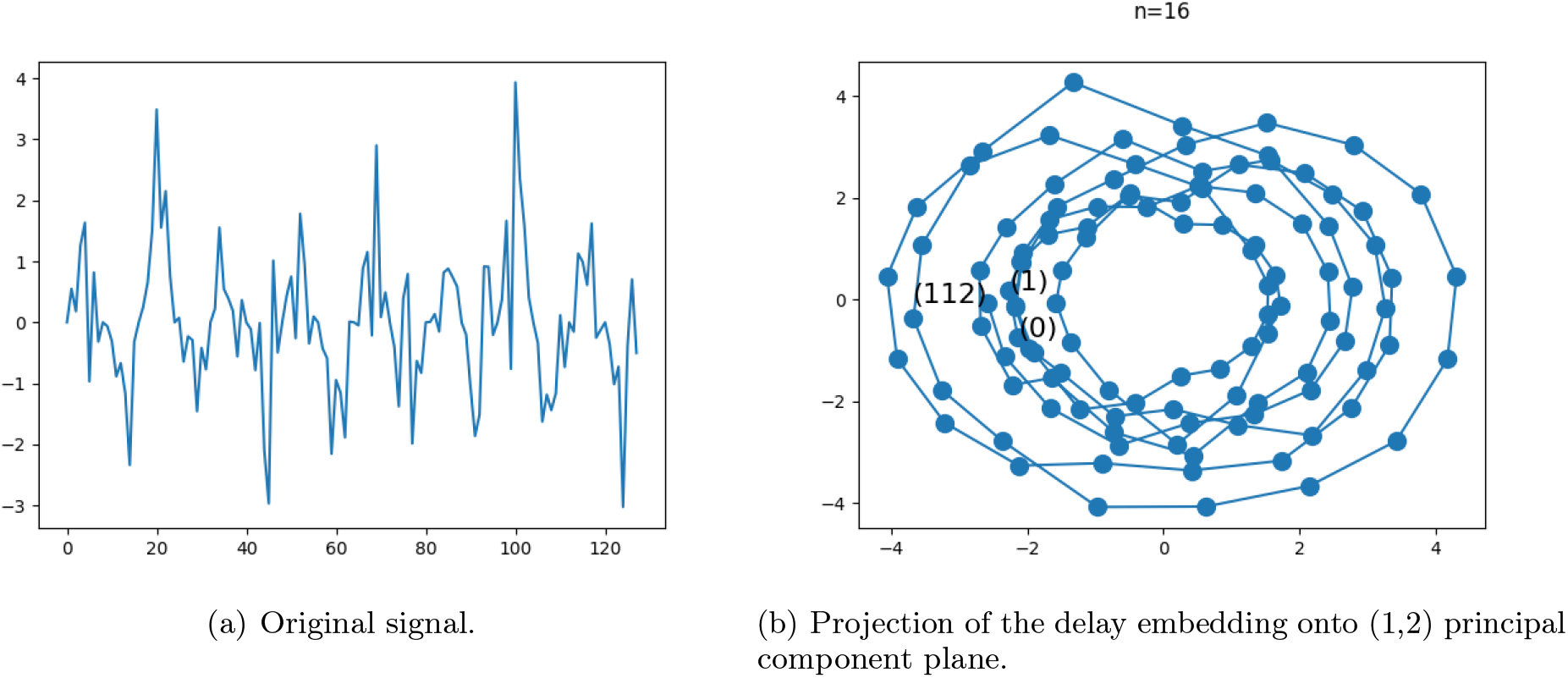
Harmonic signal with random amplitude perturbation: sin 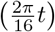, Gaussian noise (*E* = 0, *D* = 1). Parameters: *n* = 16, Δ*t* = 1, *t*_1_ = 0, *N* = 128.

This behavior confirms that noise does not alter the topological class of the signal, but only induces bounded geometric perturbations, preserving class separability in the embedding space.

### Methodological remark

In discrete Fourier analysis, the primary parameter is the signal length *N* . In contrast, the delay embedding method introduces an additional parameter, the embedding window *n*. The ratio between the minimal frequency of the signal and the window size *n* plays a crucial role in the analysis.

Importantly, the delay embedding method allows one to recover the meaningful structure of the signal in the presence of Gaussian noise even in cases where Fourier analysis does not provide unambiguous results. See Fig. 15.

**Fig. 15:**
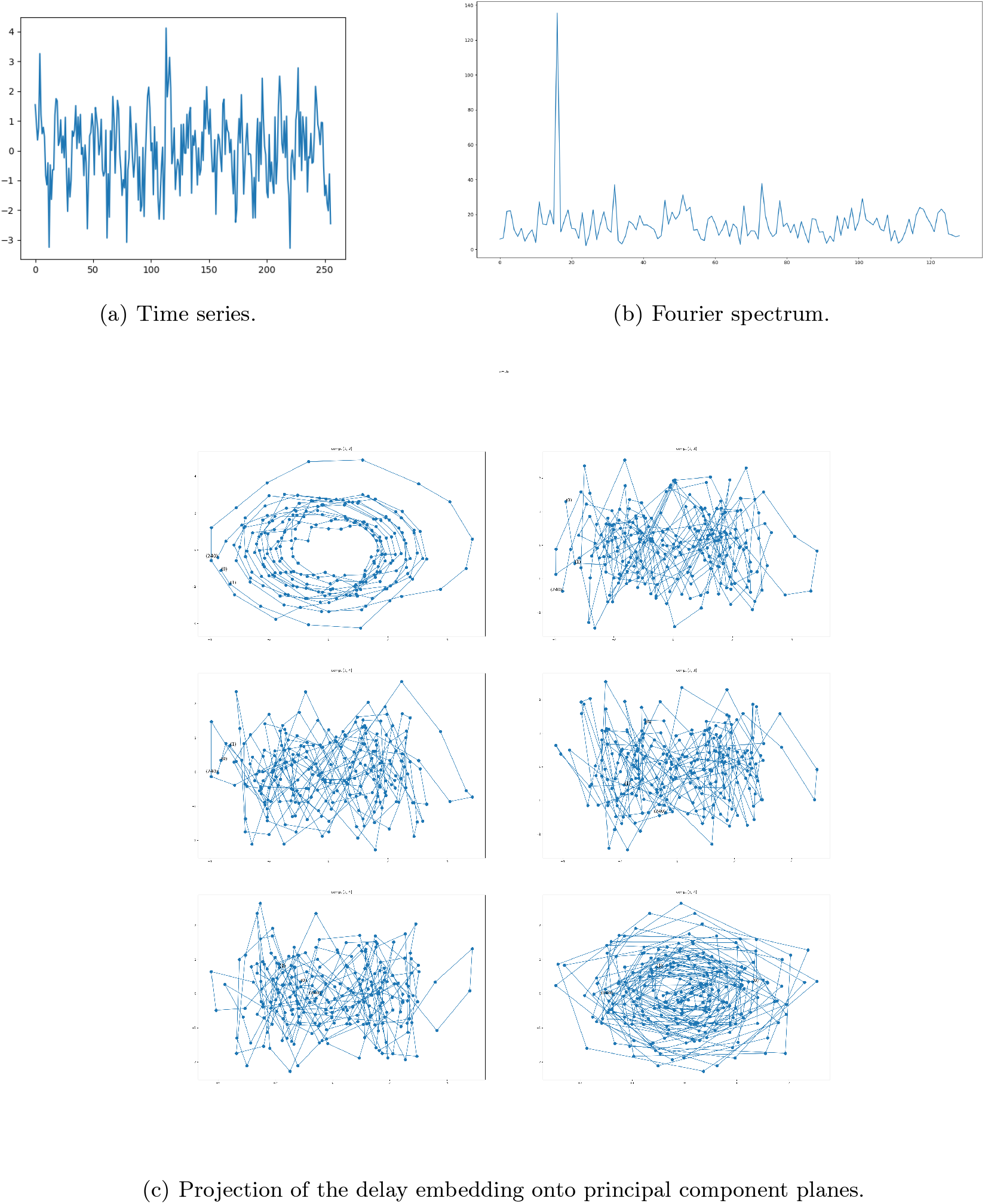
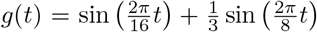 with additive Gaussian noise (*D* = 1, *E* = 0). Parameters: n = 16, Δ*t* = 1, *t*_1_ = 0, *N* = 25

### Topological classifier

#### Evaluation on synthetic data

We first demonstrate the effectiveness of the proposed algorithm on synthetic time series representing complex, noisy, multi-component signals. The constructed signals consist of superpositions of exponential, harmonic, and exponentially modulated oscillatory components, combined with additive Gaussian noise and additional perturbations such as random phase and amplitude variations.

Despite the presence of strong noise and overlapping components, the delay embedding preserves the underlying geometric structure of the signal. In particular, the embeddings exhibit low-dimensional subspaces and characteristic projection patterns corresponding to the individual components, enabling their identification.

These results confirm that the proposed topological classifier is capable of reliably recovering the intrinsic structure of complex signals (*g*_*class*1_(*t*) = *e*^−0.001*t*^ sin 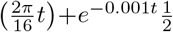 sin 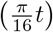, *g*_*class*2_(*t*) = *e*^−0.003*t*^ sin 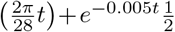 sin 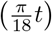 with additive Gaussian noise: *D* = 0.1, *E* = 0) and distinguishing between different generative mechanisms even in challenging noisy settings (see Fig. 16).

**Fig. 16:**
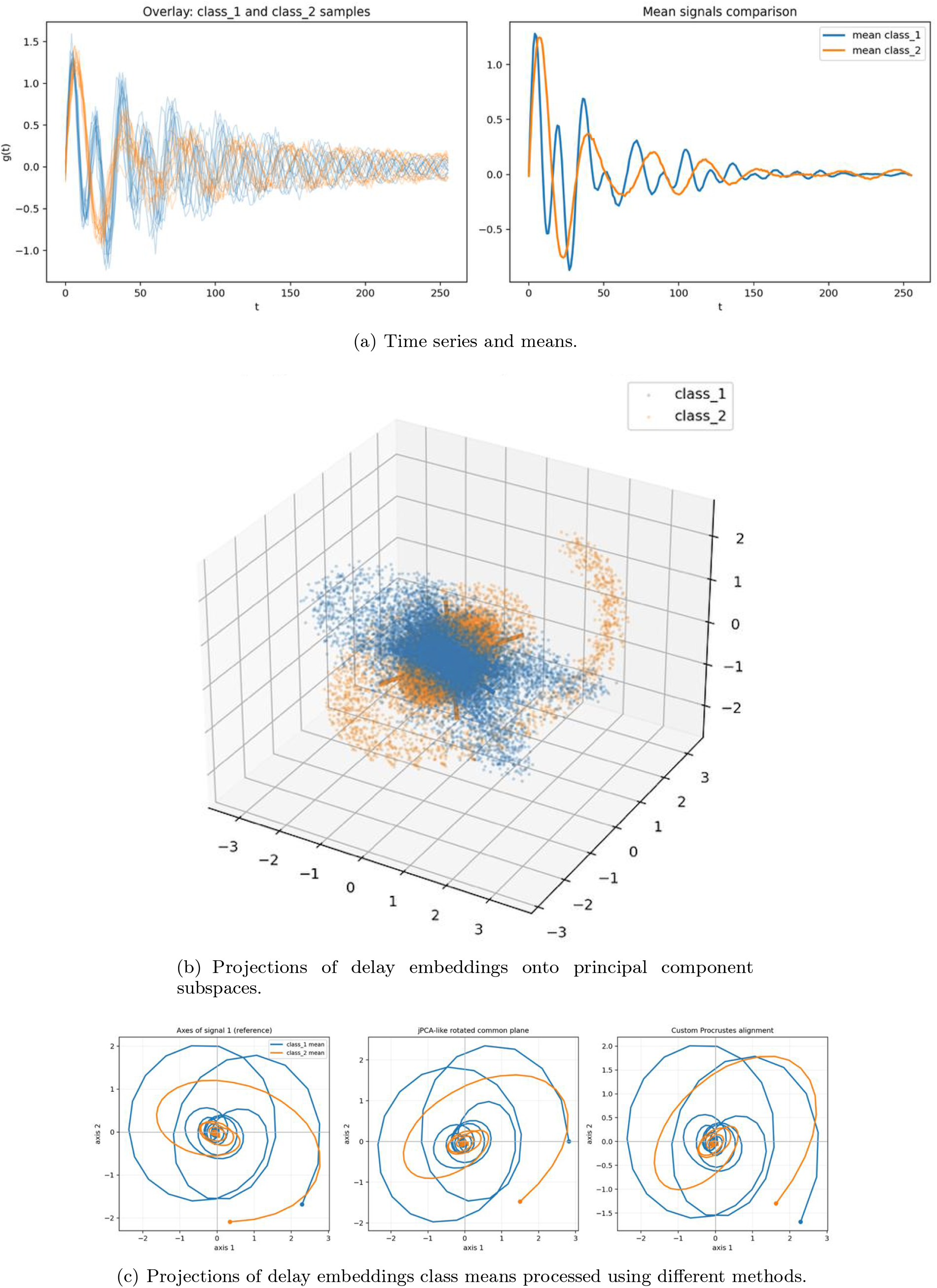
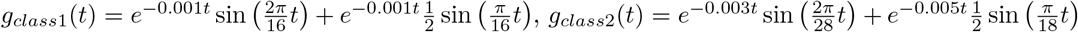 with additive Gaussian noise (*D* = 0.1, *E* = 0). Parameters: *n* = 16, Δ*t* = 1, *t*_1_ = 0, *N* = 256.

#### Evaluation on real-world data

The experiments were conducted on the publicly available EEG dataset from the UCI repository, widely used for benchmarking classification methods in biomedical signal processing.

The proposed topological classifier achieves performance that is competitive with, and in several cases exceeds, existing state-of-the-art approaches for time series classification. In particular, the obtained accuracy surpasses classical machine learning methods based on spectral features, wavelet representations, and kernel classifiers, which typically operate in the range reported in prior studies.

This improvement can be attributed to the ability of the proposed method to capture intrinsic geometric structure of the signal in the embedding space, rather than relying on pointwise or frequency-domain representations. By exploiting the topology of delay embeddings, the classifier effectively distinguishes between signals generated by different dynamical mechanisms.

These results indicate that topological analysis of embeddings provides a powerful and principled alternative to conventional approaches, establishing the proposed method as a competitive state-of-the-art solution for structured time series classification. (see Table 1 and Fig. 17). The proposed topological classifier outperforms classical machine learning and hybrid approaches while requiring no feature engineering or training.

**Table 1:** Comparison of classification accuracy with existing methods. * – mean *±* std over 10 runs. These results indicate that the proposed topological classifier achieves state-of-the-art performance on this benchmark, outperforming classical machine learning and hybrid approaches while requiring no feature engineering or training.

| Source | Classifier | Accuracy (%) |
| --- | --- | --- |
| Ekaputri et al. [21] | SVM | 77.80 |
| Ehlers et al. [22] | Discriminant analysis | 88.00 |
| Rieg et al. [23] | Random forest | 96.67 |
| Sharma et al. [24] | LS-SVM | 97.08 |
| Manivannan et al. [25] | LASSO+BDA+EANN | 99.59 |
| This work | Topological Classifier | 99.87 ± 0.1 * |

**Fig. 17:**
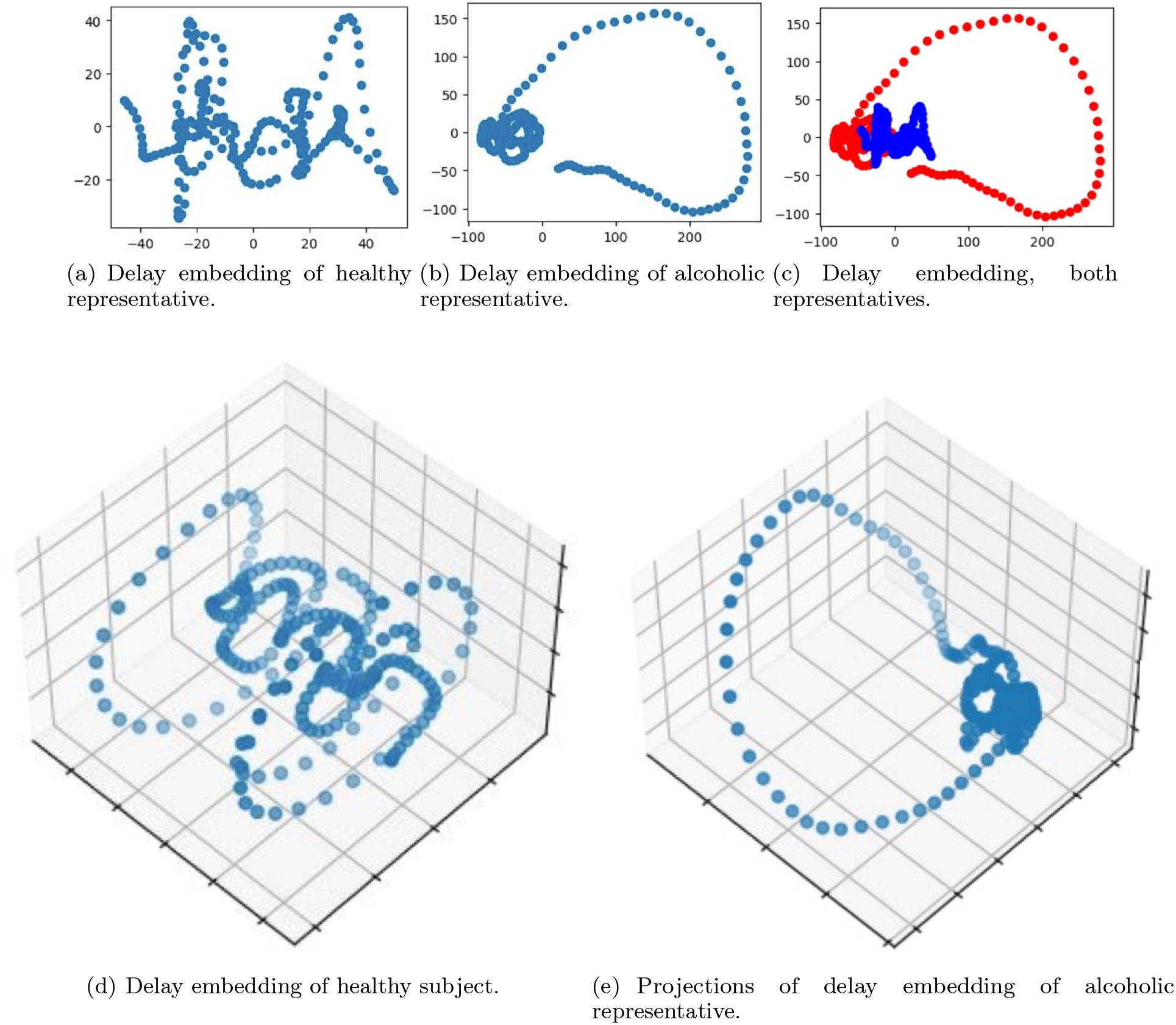
Projections of delay embeddings onto principal component subspaces reveal distinct geometric structure for time series corresponding to alcoholic and control subjects.

#### Interactive demonstration

To facilitate reproducibility and provide an intuitive understanding of the proposed framework, we developed an interactive web-based demonstration of the topological classifier. The interface allows users to construct synthetic time series by specifying combinations of exponential, harmonic, and exponentially modulated oscillatory components, and to observe the corresponding delay embeddings and their projections onto principal subspaces.

Users can explore how variations in parameters affect the geometry of embeddings, including transitions between linear, planar, and spiral structures, as well as the impact of noise through the formation of *ε*-neighborhoods. This interactive environment highlights the direct relationship between generative mechanisms and embedding geometry, and provides a visual interpretation of the classification process.

The demonstration is available at: https://data.aicumene.com.

#### Summary of results

The experiments demonstrate that:

- time series embeddings exhibit class-specific low-dimensional structure,
- these structures are well separated in the Grassmann space,
- classification based on subspace proximity is both accurate and robust,
- the proposed approach outperforms spectral methods in structurally ambiguous cases.

On a benchmark EEG dataset, the proposed topological classifier achieves performance comparable to or exceeding state-of-the-art methods. Importantly, this improvement is obtained without relying on large-scale training or feature engineering, and instead emerges from the explicit representation of the underlying dynamical structure.

These results suggest that the observed performance gain is not merely incremental, but reflects a fundamental advantage of geometric representations over spectral and feature-based approaches in structurally ambiguous settings.

The results confirm that classification is determined by the geometry of embeddings rather than pointwise signal features or spectral coefficients.

Signals generated by different dynamical mechanisms occupy distinct regions of the Grassmann manifold, which enables reliable classification even when traditional methods fail.

## Discussion

The results presented in this work suggest a fundamental shift in the way time series should be represented and analyzed. Rather than treating signals as sequences of observations or collections of spectral components, we show that they can be understood as geometric objects which structure directly reflects the underlying dynamics that generate them.

A central implication of this perspective is that the intrinsic dimension and geometry of delay embeddings act as invariants of the generative process. In contrast to classical representations, which are often sensitive to parameterization and observation scale, the embedding geometry encodes structural information that is preserved across transformations, sampling schemes, and moderate perturbations. This allows signals produced by fundamentally different mechanisms to be distinguished even when their spectral or statistical characteristics are nearly identical.

From this viewpoint, time series classification can be reformulated as a problem of geometric inference. Instead of operating in the space of signal values or engineered features, classification reduces to identifying the subspace (and its associated *ε*-neighborhood) in which the embedded trajectory resides. In this sense, the problem is naturally expressed on a Grassmann manifold, where each class corresponds to an equivalence class of subspaces.

This reformulation provides a principled explanation for the robustness of the proposed method. Noise does not destroy the structure of the signal, but induces a controlled deformation of its embedding, resulting in an *ε*-neighborhood around the underlying subspace. Classification remains stable as long as these neighborhoods do not overlap, linking statistical robustness to geometric separation.

While deep learning models may achieve high predictive performance, they typically require large-scale labeled datasets and do not provide explicit access to the underlying dynamical structure, in contrast to the proposed geometric framework.

The proposed framework also clarifies the limitations of widely used approaches. Spectral methods, including Fourier analysis, represent signals in terms of basis expansions but do not capture the generative structure of the dynamics, leading to ambiguity between signals with similar frequency content. Feature-based and machine learning approaches, while often effective, rely on representations that are not intrinsically tied to the dynamics and therefore require extensive training or domain-specific engineering. In contrast, the geometric representation developed here is directly induced by the signal-generating mechanism and does not depend on external feature construction. This suggests that the problem of time series classification is inherently geometric rather than statistical.

Beyond its immediate application to classification, the framework establishes a broader connection between time series analysis, dynamical systems theory, and geometric learning. Delay embeddings, traditionally used for phase space reconstruction, are reinterpreted here as mappings into a space where dynamical complexity becomes geometrically observable. In this sense, the intrinsic dimension of the embedding can be viewed as a measure of dynamical complexity, while the arrangement of subspaces encodes relationships between different classes of processes.

The current framework is primarily applicable to signals that can be approximated by finite superpositions of structured components. Extension to highly non-stationary or chaotic systems remains an important direction for future research.

Several limitations and open questions remain. The choice of embedding window *n* plays a crucial role in resolving the geometric structure, and while theoretical and empirical guidelines can be provided, a fully adaptive selection strategy remains to be developed. Furthermore, the current framework focuses on signals that can be approximated by finite superpositions of structured components; extending these results to non-stationary, chaotic, or high-dimensional systems represents an important direction for future work.

Finally, the geometric formulation introduced here opens the possibility of integrating invariant representations with learning-based methods. Hybrid approaches combining subspace geometry with statistical learning or neural architectures may provide a path toward scalable and interpretable models capable of handling large and heterogeneous datasets.

Taken together, these results suggest a shift from statistical and feature-based descriptions of time series toward a geometric theory in which structure, complexity, and classification are governed by the topology of embeddings. This perspective provides a unifying framework for understanding time series and lays the foundation for future developments at the intersection of dynamical systems, topology, and machine learning. This suggests that geometry, rather than statistics, may constitute the natural language of time series analysis.

## Conclusion

We have introduced a geometric framework for time series analysis that establishes a direct correspondence between the generative structure of a signal and the topology of its delay embedding. We show that broad classes of signals (including exponential, harmonic, and exponentially modulated oscillatory processes) induce low-dimensional subspaces which dimension reflects the number and type of latent dynamical components.

Building on this result, we reformulate time series classification as the identification of subspaces together with their associated *ε*-neighborhoods. This leads to a topological classifier that is interpretable, data-efficient, and robust to noise, where perturbations correspond to bounded geometric deviations in embedding space.

More broadly, these results suggest that the intrinsic dimension and geometry of delay embeddings act as invariants of the underlying dynamics. This shifts the perspective on time series analysis from signal-based and feature-based representations toward a geometric description in which structure and complexity are directly observable.

Future work will focus on extending the framework to non-stationary and high-dimensional signals, developing adaptive strategies for embedding selection, and integrating geometric representations with learning-based models.

An interactive web-based demonstration of the proposed framework is available, providing an experimental environment for exploring the geometric signatures of dynamical systems through synthetic signal construction and visualization.

Overall, the proposed approach lays the foundation for a geometric theory of time series, in which classification, representation, and dynamical structure are unified through the topology of embeddings.

In this sense, the proposed framework points toward a unified geometric theory of time series, where dynamics, structure, and inference are governed by topology rather than statistics.

## Acknowledgements

This work was supported by Neurosputnik LLC. The authors thank colleagues for valuable discussions that contributed to this study.

